# Strong replicators associated with open chromatin are sufficient to establish an early replicating domain

**DOI:** 10.1101/289025

**Authors:** Caroline Brossas, Sabarinadh Chilaka, Antonin Counillon, Marc Laurent, Coralie Goncalves, Bénédicte Duriez, Marie-Noëlle Prioleau

## Abstract

Vertebrate genomes replicate according to a precise temporal program strongly correlated with their organization into topologically associating domains. However, the molecular mechanisms underlying the establishment of early-replicating domains remain largely unknown. We defined two minimal *cis*-element modules containing a strong replication origin and chromatin modifier binding sites capable of shifting a targeted mid-late replicating region for earlier replication. When inserted side-by-side, these modules acted in cooperation, with similar effects on two late-replicating regions. Targeted insertions of these two modules at two chromosomal sites separated by 30 kb brought these two modules into close physical proximity and induced the formation of an early-replicating domain. Thus, combinations of strong origins and *cis*-elements capable of opening the chromatin structure are the basic units of early-replicating domains, and are absent from late-replicated regions. These findings are consistent with those of genome-wide studies mapping strong initiation sites and open chromatin marks in vertebrate genomes.

## Introduction

A precise, cell type-specific temporal program governs the duplication of vertebrate genomes (Ryba et al., 2010). Replication timing (RT) profiles consist of large regions (from 200 kb to 2 Mb) of DNA homogeneously displaying early or late replication (constant timing regions, or CTRs) punctuated by gradients of progressive change in RT, known as timing transition regions (TTR) (Hiratani et al., 2008). Large early CTRs result from the more or less synchronous activation of clusters of origins. The genome-wide mapping of replication origins with different methods has revealed a high density of efficient site-specific origins in early-replicated domains, whereas late regions are usually origin-poor (Prioleau and MacAlpine, 2016). These findings suggest that early and late replication domains are formed through different, as yet uncharacterized molecular mechanisms. RT domains are correlated with the organization of chromosomes into two main types of compartments: compartment A, which is open and replicated early, and compartment B, which is closed and replicated late (Ryba et al., 2010). These compartments, which display greater interaction within themselves rather than across them, were initially defined by the Hi-C method, at a resolution of 1 megabase (Lieberman-Aiden et al., 2009). A comparison of higher resolution Hi-C data defining submegabase domains called TADs (topologically associating domains) with RT domains, revealed a correlation of their boundaries (Pope et al., 2014). This observation led to a model (the replication domain model) in which each RT domain acts as a regulatory unit, determining when the replicons within its boundaries can fire. The zinc finger protein CTCF found enriched at TADs boundaries, has been implicated in the local insulation of chromosome neighborhoods through its enhancer-blocker activity (Bell et al., 1999; Dixon et al., 2012; Hou et al., 2010; Sexton et al., 2012). However, CTCF depletion does not disrupt the organization of chromosomes into A/B compartments, revealing that local insulation and higher-order compartmentalization are based on different molecular determinants (Nora et al., 2017). The direct molecular drivers of CTCF-independent higher-order compartmentalization remain to be defined, but are probably also involved in RT control.

The only major player in the RT program for which the mode of action has been elucidated is Rif1 (Cornacchia et al., 2012; Yamazaki et al., 2012). Rif1 has a global repressive effect on genome-wide DNA replication. This effect is mediated by the recruitment of protein phosphatase 1, which opposes the Dbf4-dependent kinase (DDK) activity required for origin firing (Hiraga et al., 2014). Consistent with this direct role, ChIp analyses of Rif1 throughout the mouse genome revealed an overlap between Rif-1 associated domains and late replication (Foti et al., 2016). However, only regions associated with Rif1 but not with the nuclear lamina switch to early replication in a context of Rif1 depletion. This finding suggests that other major players, such as the nuclear lamina, are also involved in controlling late replication. One key question that remains to be resolved concerns whether late-replicating domains are so robustly constrained by their nuclear compartmentalization that the targeting of a very efficient origin associated with early timing control elements could not locally advance the timing of their replication. The underlying question is whether a late domain is defined by the deficiency of an early firing signal together with an accumulation of signals imposing the late firing of many potential initiation sites. A related question is whether early-replicated domains are defined solely by the absence of a strong negative signal, such as association with the nuclear lamina and/or Rif 1. Alternatively, early CTRs may result fortuitously from the more or less synchronous firing of a cluster of replicons, each with its own individual local early timing elements. This hypothesis is supported by the high density of efficient origins proximal to sites associated with open chromatin marks in early-replicated domains (Picard et al., 2014). We previously tested this hypothesis by determining whether it was possible to construct *de novo* an autonomously regulated domain of replication inside a naturally mid-late-replicating domain. We investigated key timing control *cis*-elements in avian DT40 cells, by creating several constructs targeting a specific locus replicated in mid-late S-phase and assessing their impact on DNA RT (Hassan-Zadeh et al., 2012). We identified a minimal combination of *cis*-regulatory elements capable of inducing a significant shift in RT to earlier time points. This minimal combination of DNA elements consists of a strong origin (the *β^A^ globin* promoter) flanked by two copies of binding sites for the USF (upstream stimulating factor, FIV of the chicken HS4 insulator) transcription factor. We found that both a strong replicator and flanking USF binding sites were required for a shift in RT. However, the origin function of the *β^A^-globin* promoter was not dependent on the presence of *cis*-elements controlling RT, making it possible to separate origin function from timing information (Valton et al., 2014). We showed that USF binding sites can direct specific histone modifications, as the chromatin landscape of the advanced origin displays a switch from histone marks characteristics of closed chromatin (H3K27me3) to more open chromatin marks (acetylated histone H3, H2AZ and dimethylated H3K4), consistent with the previously reported function of USF in preventing the encroachment of heterochromatin (West et al., 2004). This observation highlights the local role of histone modifications in determining the RT of an adjacent replication origin (Goren et al., 2008). We also showed that the impact of this group of elements on RT could be increased further by the presence of a gene under the control of an active strong constitutive *β-actin* promoter (Hassan-Zadeh et al., 2012). We show here that the cooperation of two closely spaced replicators is associated with the formation of a 5.6 kb domain in which the chromatin is more accessible. This small domain can counteract the potential constraints imposed by late-replicating domains. Moreover, when inserted into a mid-late-replicating region at two positions 30 kb apart, this combination can generate an early-replicating domain and bring the two modules into close spatial proximity. Overall, our synthetic approach sheds new light on the mechanisms responsible for the establishment and maintenance of early- and late-replicating domains.

## Results

### Quantitative analysis of the impact of *cis-*regulatory elements on replication timing reveals cooperation between two minimal autonomous replicons

We previously investigated the impact of *cis*-regulatory elements on replication timing (RT), by developing a method for precise quantification of the magnitude of the shift in RT induced by the insertion of an ectopic DNA sequence into a specific mid-late-replicating locus ((Hassan-Zadeh et al., 2012) and Figure 1). In this study, we analyzed a much larger number of cell lines than previously described to quantitatively evaluate the variability of RT shifts imposed by our different constructs in the same mid-late replicated region (chr1:74,813,240 Assembly WUGSC 2.1/galGal3 May 2006, chr1:72,565,520 Assembly Gallus_gallus-5.0/galGal5 Dec 2015) (Table S1). The calculation of ΔL and ΔE values involves the conversion of the RT shift into a difference in NS (newly synthesized strand) enrichment relative to an internal reference (without, unmodified allele) specific to each experiment and, thus, independent of slight experimental variations between clones during cell sorting. The results obtained with this method therefore provide an accurate estimate of the extent of the shift in RT (Figure 1B).

**Figure 1:**
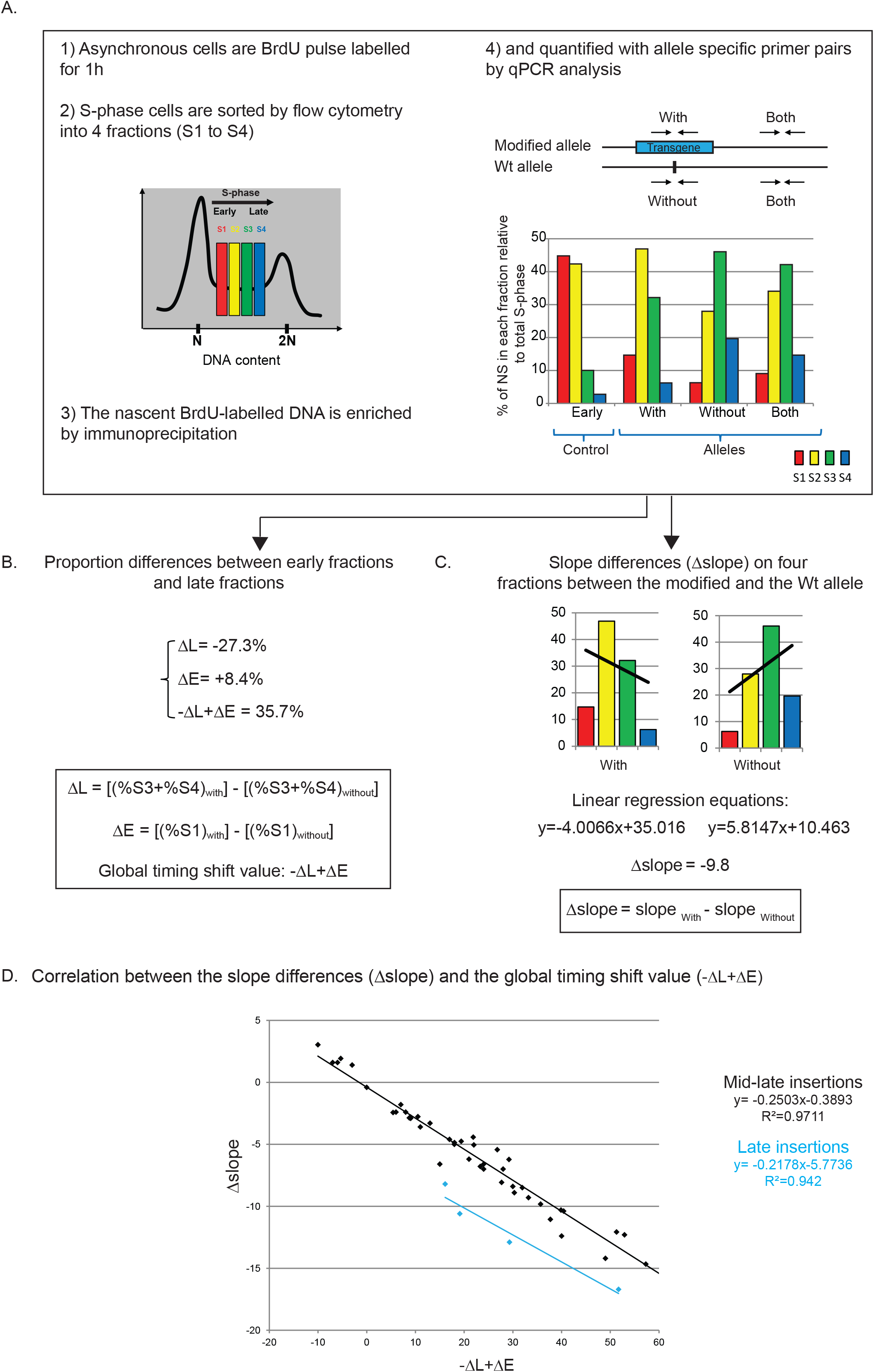
Methods for allele-specific analysis of replication timing by real-time PCR quantification. (A) BrdU pulse-labeled cells were sorted into four S-phase fractions (S1 to S4) and the immunoprecipitated newly synthesized strands (NS) were quantified by real-time qPCR in each fraction. Specific primer pairs determine the replication timing (RT) profile for the modified allele (With), the wt allele (Without) and both alleles (Both). The endogenous *β-globin* locus was analyzed as an early-replicated control (Early). (B and C) Two methods of calculation were used to determine the difference in RT between the wt and modified alleles: the -ΔL+ΔE method (B) and the Δslope method (C). (D) Analysis of the correlation between the Δslope method and the -ΔL+ΔE method for all clonal cell lines previously published (Hassan-Zadeh et al., 2012; Valton et al., 2014) or analyzed for the first time here. The linear regression curve is shown, with the corresponding equation and the coefficient of determination (R^2^).

We quantitatively compared the RT shifts triggered by different constructs by plotting the distributions of timing shift values (-ΔL+ΔE) for each transgene and compared them with a reference corresponding to cell lines containing a non-shifting construct lacking a replicator (Figure 2A (i) and (Valton et al., 2014)). For eight clones (2xFIV line), we confirmed that the *β^A^-globin* promoter containing a strongly active replication origin ((Hassan-Zadeh et al., 2012) and (Valton et al., 2014)) flanked by USF binding sites significantly advanced RT at position of the insertion (Figure 2A (ii), *p*-value =4.57E-05). Moreover, an analysis of six cell lines containing the active *β-actin* promoter together with an active replication origin ((Hassan-Zadeh et al., 2012) and Figure S1A) (*BsR*) confirmed the ability of this promoter to advance RT significantly (Figure 2A (iii), *p*-value = 3.38E-03). Similar shifts were observed for the 2xFIV clones and the *BsR* clones (Figure 2A, *p*-value = 1.00 between (ii) and (iii)), although the - ΔL+ΔE values were more dispersed for *BsR* clones (interquartile range (Q1-Q3) of 5.73 for 2xFIV and 11.4 for *BsR* distributions, respectively). We then assessed the impact on RT of the combination of these two minimal modules in eight clonal cell lines. Our results confirmed that the large 2xFIV+*BsR* construct imposed a stronger shift to earlier replication at the inserted locus than the presence of a single minimal module alone (Figure 2A (iv), *p*-value = 3.25 E-03 between (ii) and (iv); *p*-value = 5.93E-02 between (iii) and (iv) and Figure S1B, S2B and C and (Hassan-Zadeh et al., 2012)).

**Figure 2:**
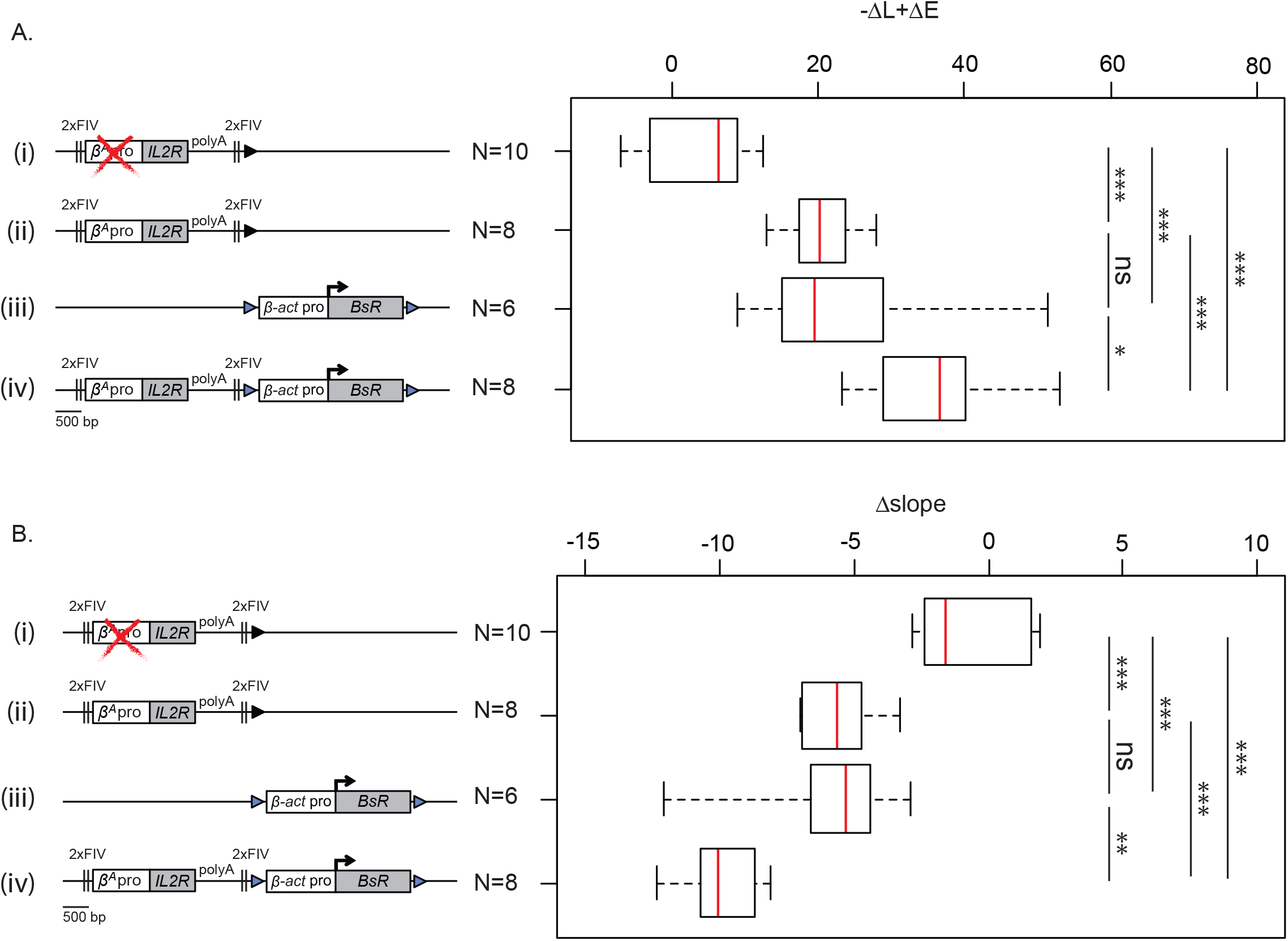
Quantitative analysis of replication timing shifts reveals cooperation between two combinations of *cis*-regulatory elements. (A and B) Distributions of RT shifts calculated with the -ΔL+ΔE method (A) or with the Δslope method (B). Different transgenes in a mid-late-replicating region are compared: the *IL2R* reporter gene under the control of the *β^A^-globin* promoter (*β^A^* pro) containing an inactive (i) or an active origin (ii) flanked by two copies of the USF-binding sites (2xFIV), the *blasticidin* resistance gene (*BsR*) under the control of the *β-actin* promoter (*β-act* pro) (iii) or a combination of these two transgenes (2xFIV+*BsR*) (iv). Rectangle edges correspond to the 0.25 and 0.75 quartiles, the red lines represent the median and the whiskers extend to the smallest and largest (-ΔL+ΔE or Δslope) values. Statistical analysis was performed with Wilcoxon nonparametric two-tailed tests (ns, not significant; \**p*<0.1; \*\**p*<0.05; \*\*\**p*<0.01).

We developed a second method to confirm the results of our quantitative analysis (Figure 1C). We first plotted the trend curves fitting the NS enrichment distributions quantified in the four S-phase fractions for the modified (With) and wt allele (Without). The global rate of change in NS enrichment between the two alleles was then determined by calculating the difference in slopes of regression lines for the wt and modified alleles. Similar significant shifts were detected after the insertions of the 2xFIV and *BsR* minimal modules (Figure 2B, *p*-value = 4.4E-04 between (i) and (ii); *p*-value = 1.34E-03 between (i) and (iii), *p*-value = 0.60 between (ii) and (iii)) and a stronger significant shift was observed when the two minimal modules were inserted together (2xFIV+*BsR*) (Figure 2B, *p*-value = 9.31E-04 between (ii) and (iv); *p*-value = 3.30E-02 between (iii) and (iv)). An analysis of the correlation between the results of the two methods based on published RT values for cell clones and the values obtained for the new clones included in this study demonstrated that the two methods are comparable and accurate for calculating the magnitude of the RT shift in the mid-late region (Figure 1D, R^2^ =0.97, black regression line).

### The two small modules responsible for the RT shift have different chromatin architectures

It has been suggested that chromatin architecture makes a major contribution to RT control (Rhind and Gilbert, 2013). We analyzed the 2xFIV and *BsR* shifting modules inserted separately or in combination (2xFIV+*BsR*). The heterochromatin region located upstream from the early-replicating *β-globin* locus was used as a control for the condensed state (Litt et al., 2001; Prioleau et al., 1999). For all cell lines, we detected similar small amounts of DNA release from this region, for all MNAse concentrations (Figures 3A-C, cond1 and 2). We detected similar behavior at the insertion site in the wt allele and 5 kb downstream from this site (insertion site +5 kb), indicating that the targeted genomic region is not naturally accessible, consistent with the global genomic landscape around the insertion site, with an absence of actively transcribed genes. By contrast, the transcriptionally active *MED14* promoter (Figure 3E) displayed a higher rate of nucleosome release (Figure 3 A-C). At low MNAse concentration (2.5 U), the nucleosomes embedded within the entire 2xFIV construct were released similarly to those on the wt allele, indicating that the 2xFIV construct has no major overall impact on chromatin accessibility. However, at the *β^A^-globin* origin site (*β^A^* promoter 2), nucleosome release rates were higher for the highest MNAse concentration (160 U), whereas the upstream tiled amplicon (*β^A^* promoter 1) behaved in the same way as the condensed *β-globin* region (Figure 3A). This result suggests that there is a nucleosome positioned at this site, protecting the chromatin from MNAse digestion. This finding is consistent with those of a previous study reporting strong nucleosome positioning sites around the TSS of the endogenous chicken *β^A^-globin* gene (Davey et al., 1995). We also found that nucleosomes embedded within the 2xFIV elements at both upstream and downstream positions (5’ 2xFIV and 3’ 2xFIV) become more susceptible to MNAse digestion at higher concentrations, indicating higher levels of chromatin exposure at these sites, possibly due to the capacity of these elements to recruit histone modifiers upon USF binding (West et al., 2004 and Figure 3D). Overall, these results indicate that the 2xFIV construct is bounded by two sites of moderately accessible chromatin surrounding a strong replicator composed of a well-positioned nucleosome. By contrast, for the *BsR* construct, nucleosome release levels were 10 times higher than for the wt chromosome and condensed control regions at low MNAse concentration (Figure 3B). We concluded that the active *β-actin* promoter containing a strong replicator imposes an accessible chromatin structure on the promoter and the transcribed region.

**Figure 3:**
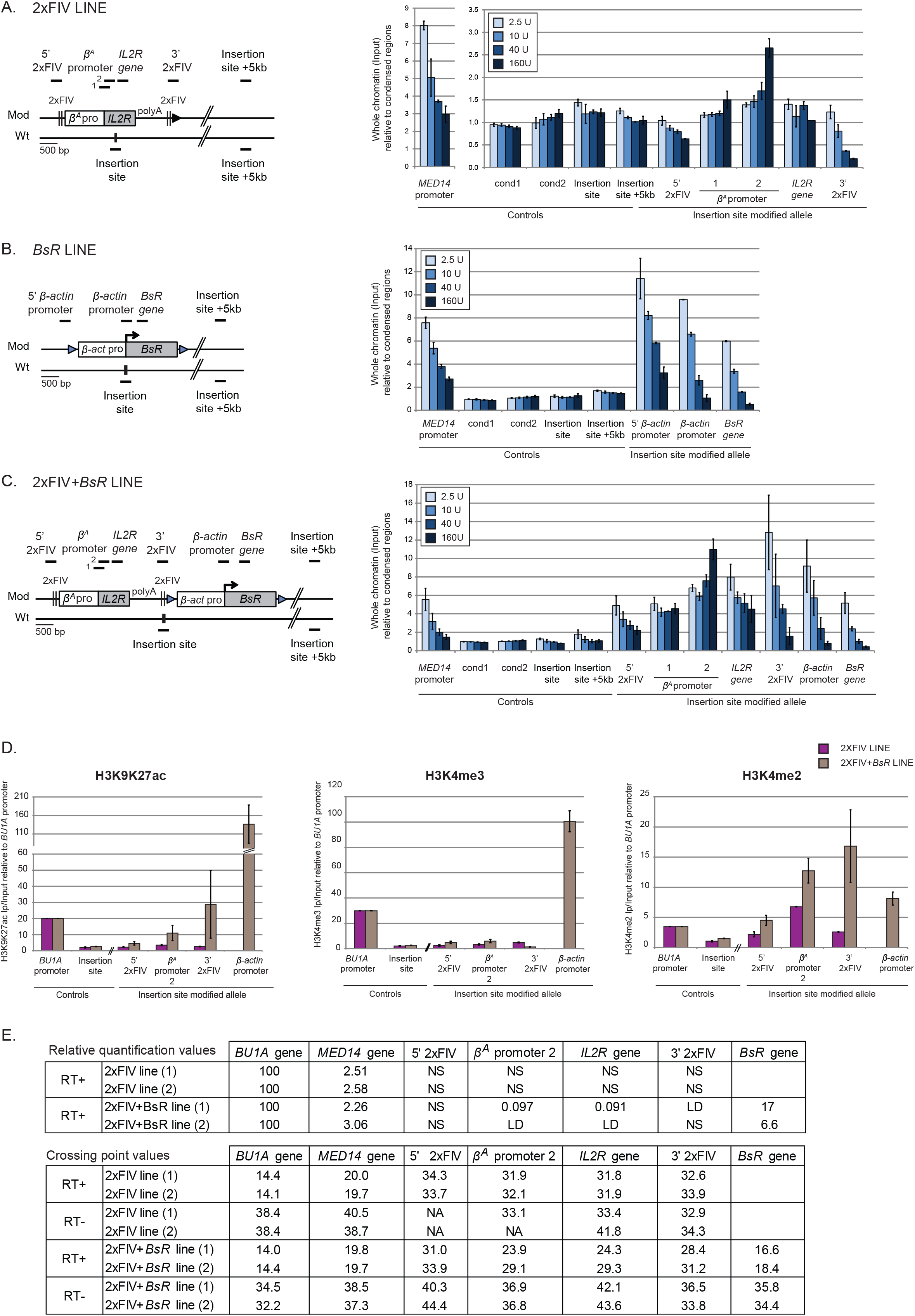
Different chromatin architectures are associated with each combination of *cis*-regulatory elements. The 2xFIV (A), *BsR* (B) and 2xFIV+*BsR* (C) transgenes are shown on the left, with the positions of the amplicons used for quantification (thick black lines with names indicated above). The endogenous active *MED14* promoter or the endogenous active *BU1A* promoter, two genomic regions located within the condensed region upstream from the *β-globin* locus (cond1 and cond2) were analyzed as controls. Quantification, by real-time qPCR, of total chromatin (input) extracted from two clones of the 2xFIV (A), *BsR* (B) and 2xFIV+*BsR* (C) cell lines after digestion with increasing concentrations of micrococcal nuclease (MNAse, 2.5U, 10U, 40U, 160U/mL). (D) Immunoprecipitations of H3K9K27 acetylation (ac), H3K4 trimethylation (me3) or H3K4 dimethylation (me2) on formaldehyde-crosslinked chromatin extracted from the 2xFIV (purple) and 2xFIV+*BsR* (gray) cell lines. Data are presented as enrichments of immunoprecipitated material relative to input DNA and normalized against *BU1A* enrichment. (E) Relative quantification, by real-time qPCR, of mRNA expression levels (RT-PCR+) or background levels (RT-PCR-), was performed in the 2xFIV and 2xFIV+*BsR* cell lines. For relative quantification, mRNA levels were normalized against *BU1A* mRNA levels arbitrarily set at 100 (first table). Crossing point (Cp) values for RT-PCR+ and RT-PCR-experiments are reported in the second table. NA corresponds to non-amplified samples, NS to non-specific signals and LD to the limit of detection. Error bars indicate the standard deviation for qPCR triplicates (A, B and C) or duplicates (D, E) for two independent clones.

### The cooperation of the two modules in RT control is associated with a propagation of open chromatin structure

We then investigated the organization of the 2xFIV+*BsR* construct displaying the strongest replication shift. The 2xFIV+*BsR* construct induced a large increase in nucleosome release for the 2xFIV transgene at low MNAse concentration (2.5 U) (approximatively five times more DNA released than for the 2xFIV construct alone; compare Figures 3A and C). By contrast, combination yielded a similar pattern of accessibility to the *BsR* transgene alone; compare Figures 3B and C. We then characterized specific histone marks at several positions within the 2xFIV and 2xFIV+*BsR* transgenes (Figure 3 D). The *BU1A* promoter was used as a positive control (Sarkies et al., 2012). The insertion site has no active histone marks, such as H3K4 methylation or H3K9K27 acetylation, consistent with its profile of poor chromatin accessibility (insertion site, Figure 3D). By contrast, within the 2xFIV+*BsR* transgene, the *β-actin* promoter recruited high levels of active H3K9K27ac and H3K4me3 marks, together with moderate levels of H3K4me2 marks. The H3K4me3 mark was not detected at any other position within this construct or within the 2xFIV construct. The density of H3K9K27ac marks gradually decreased from the *β-actin* promoter to the upstream part of the 2xFIV+*BsR* construct. Finally, within the 2xFIV+*BsR* transgene, the H3K4me2 mark was strongly recruited at the 3’ 2xFIV element, its density then gradually decreasing in the upstream part of the transgene. We also showed that the 2xFIV module alone was covered by moderate levels of H3K4me2 marks on the 5’ and 3’ 2xFIV elements, associated with a stronger enrichment on the *β^A^-globin* promoter. This result is consistent with previous observations for chromatin accessibility indicating that the 2xFIV module is located in a slightly more accessible chromatin environment. Overall, the deposition of histone marks correlates well with chromatin accessibility. We then quantified mRNA levels at different positions within the transgenes, by RT-qPCR in two clones of 2XFIV and 2xFIV+*BsR* cell lines (Figure 3E). *BU1A* mRNA level was set arbitrarily to 100 as a reference. The amount of *MED14* mRNA confirmed that this locus was actively transcribed in DT40 cells. For the 2xFIV construct, we detected no significant transcription for the *IL2R* reporter gene (Figure 3E; 2xFIV line; *β^A^* promoter 2 and *IL2R* gene) or at several other positions within the transgene. The *β^A^-globin* promoter/origin retained its origin function, but its erythroid-specific promoter function was lost in DT40 lymphoid cells. The 2xFIV RT shift module therefore carries timing information that is independent of active transcription. With the 2xFIV+*BsR* transgene, we detected transcription of the *BsR* gene in both clones analyzed (Figure 3E; 2xFIV+*BsR* line; *BsR* gene), and low levels of transcription for the *IL2R* gene (Figure 3E; 2xFIV+*BsR* line) in one cell line. Overall, these results suggest that close proximity to the highly accessible *BsR* module influences chromatin structure within the upstream 2xFIV module, leading to a shift from a slightly accessible structure to a more open state in which the two replicators (*β^A^-globin* and *β-actin)* are in an accessible environment. This 5.6 kb domain is sufficient to induce a pronounced shift in RT toward earlier replication following its insertion into a naturally closed mid-late-replicating region.

### Two late-replicating environments are permissive to a shift toward earlier replication after the site-specific insertion of a combination of timing control elements and two strong origins

It remains unknown whether late replicating regions are permissive to early replication signals. One hypothesis is that the nuclear and chromatin organizations of late-replicating domains impose robust constraints that prevent earlier firing. We addressed this question directly, by investigating whether the 2xFIV+*BsR* combination functions as efficiently when targeted to regions naturally displaying very late replication. We selected two late-replicating regions from the genome-wide RT profile obtained for wt DT40 cells. One of these regions was located 2 Mb upstream from the previously studied mid-late-replicating region on chromosome 1 (Late1 chr1:72,616,319 / chr1:70,523,649 and Mid-late chr1:74,813,240 / chr1:72,565,520 Assembly WUGSC 2.1/galGal3 May 2006 or Gallus_gallus-5.0/galGal5 Dec 2015 respectively; Figure 4A) and the other was located 105.4 Mb downstream from this mid-late-replicating region (Late2 chr1:182,214,693/chr1:177,936,192 Assembly WUGSC 2.1/galGal3 May 2006 or Gallus_gallus-5.0/galGal5 Dec 2015 respectively). We confirmed that these two regions were, indeed, replicated much later than the previously studied region (Figure 4B and C, without). We calculated both -ΔL+ΔE and Δslope values for cell clones containing the 2xFIV+*BsR* transgene inserted into late-replicating regions (Figure 4B and C) and included them in the correlation analysis performed with mid-late-replicating clones (Figure 1D). Despite the small number of clones analyzed, the -ΔL+ΔE and Δslope parameters were also found to be correlated for late-replicating clones (Figure 1D; R²=0.94). However, mid-late- and late-replicating target clones formed two distinct populations, characterized by close slopes of regression lines (Figure 1D; slope of the regression line = −0.25 for mid-late-replicating clones and −0.22 for late-replicating clones). This result suggests that it is not possible to compare the magnitude of the shift between the mid-late- and late-replicating clones by the -ΔL+ΔE method, consistent with the finding that, for late S phase-replicating loci, the signal is mostly associated with the S4 fraction for the unmodified late-replicating allele, whereas this method was initially designed for loci displaying signal enrichment mostly in S3. The new calculation method revealed a strong impact on RT of transgene insertion at the late 1 site. This shift was even stronger than that observed following insertions into the previously studied mid-late-replicating region (Δslope = −12.9 for clone 1 and −16.7 for clone 2, Figure 4B versus smallest and greatest Δslope values calculated for eight clones: −8.1 and −12.3, respectively, in the mid-late-replicating region, Figure 2B). However, in the late 2 region, we detected a shift in RT similar to that induced in a mid-late-replicating region (Δslope = −8.2 for clone 1 and −10.6 for clone 2, Figure 4C). Our findings confirm the robustness of the previously identified combination of *cis*-elements. We show here that this combination of elements can impose local autonomous control over replication timing in at least three chromosomal regions, one replicating in mid-late S phase and the other two displaying late replication.

**Figure 4:**
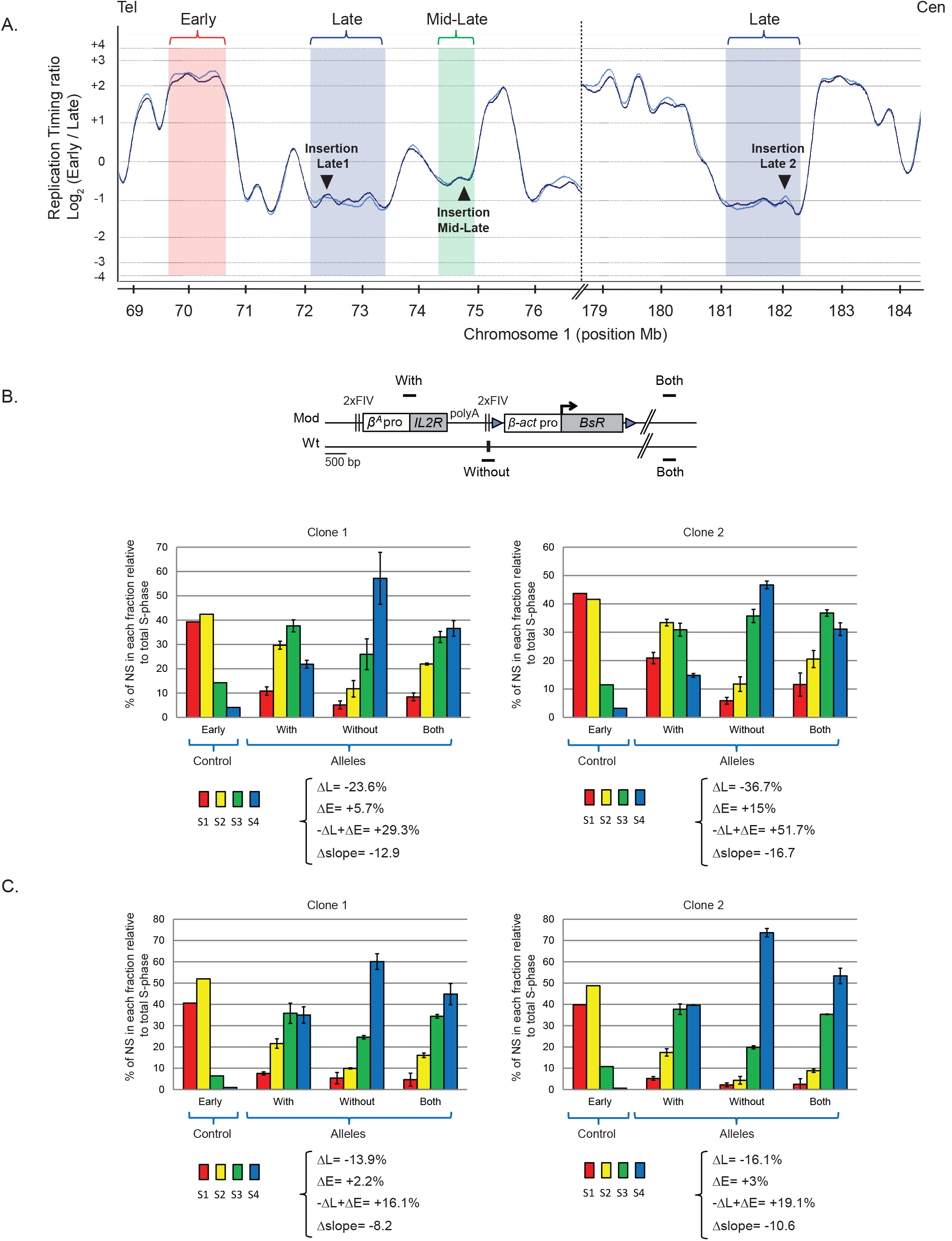
Two late-replicating environments are permissive to a shift toward earlier replication after the site-specific insertion of a combination of timing control elements and two strong origins. (A) RT profiles for two parts of chromosome 1 are shown (genomic positions: chr1:68,850,000-76,840,000; 8 Mb and chr1:178,900,000-184,500,000; 5.6 Mb; Assembly WUGSC 2.1/galGal3 May 2006). NS purified from two S-phase fractions (early and late) were differentially labeled and cohybridized onto a chicken whole-genome oligonucleotide microarray at a density of one probe every 5.6 kb. Typical early, mid-late and late replication domains are shown in red, green and blue, respectively. The sites chosen for insertion are indicated (insertion Mid-Late, insertion Late 1 and insertion Late 2). (B and C) RT profiles of chromosomal alleles following the targeted integration of a 2xFIV+*BsR* transgene into two late replicating loci, late 1 (B) and late 2 (C). The modified (Mod) and wild-type (wt) alleles are shown.

### Two advanced replicons separated by 30 kb synergize to form a synthetic early-replicating region

Early-replicating domains are characterized by the presence of strong origins or clusters of origins that can be detected by the SNS (small nascent strands) method at regular intervals of several tens of kilobases. We investigated the impact on RT of the insertion of two strong replicators carrying timing information, about 30 kb apart. We did this by inserting a second autonomous replicon 30 kb upstream of the first (insertion site 3; chr1: 72,536,061 Assembly Gallus_gallus-5.0/galGal5 Dec 2015, Figure 5A). The second autonomous replicon (2xFIV+*PuroR*) was similar to the first except that the gene used conferred puromycin-N-acetyltransferase resistance (*PuroR*). We assessed the impact of RT changes in the middle of our modified region, by introducing a reporter construct in a central position (Insertion site 2; chr1: 72,548,590 bp Assembly Gallus_gallus-5.0/galGal5 Dec 2015, Figure 5A) between the two autonomous replicons (1+2+3) or on the other chromosome (1+3) (Figure 5B). The reporter construct consisted of the erythroid-specific *β ^A^-globin* origin/promoter linked to the green fluorescent protein (GFP) reporter gene and a 1.6 kb fragment of human chromosome 7 (h.K7; chr7:26,873,165-26,874,805; Assembly (GRCh38/hg38) Human Dec. 2013) with no replication origin (Figure 5B). We found that the *GFP* reporter construct had no impact on RT when inserted alone at site 2 (-ΔL+ΔE= −5.1% and Δslope = −0.9, Figure S2A). We then showed that the 2xFIV+*BsR* construct inserted at site 1 had similar effects on RT when associated with *GFP* reporter construct insertion at site 2 on the same or the other chromosome (compare Figure S2B and C with Figure S1B and (Hassan-Zadeh et al., 2012)). These results confirm the neutral impact on RT of inserting the *GFP* reporter construct at site 2. We then investigated whether the insertion of a single autonomous replicon at site 3 was able to advance RT as efficiently as insertion at site 1. We confirmed that the 2xFIV+*PuroR* construct induced a substantial shift in RT when inserted at this new genomic position (-ΔL+ΔE= +26.8% and Δslope_f_ = −5.4 for clone 1 and -ΔL+ΔE= +22% and Δslope_f_= −5.1 for clone 2, Figure S2D). This shift was similar to that observed in 2xFIV+*BsR* clones (*p*-value = 0.2286 between (i) and (iii), Figure 5C). As previously shown, the insertion of the *GFP* reporter construct at site 2 had no impact on RT when associated with a single autonomous replicon at site 3 on either the same or the other chromosome (Figure S2, compare -ΔL+ΔE and Δslope values between D and E). We then investigated whether the insertion of two autonomous replicons at sites 1 and 3, with (1+2+3) or without (1+3) the central *GFP* reporter construct, resulted in larger shift in RT shift over the whole of the genomic region targeted. The RT of the targeted genomic region changed considerably, with a strong shift towards an even earlier replication pattern (Figure S3, -ΔL+ΔE= 63.3% and 70.3% for 1+2+3 insertions (A) and -ΔL+ΔE= 65.2%, 84.2%, 57.3%, for 1+3 insertions (B)). The shift in RT towards an earlier pattern was similar for the 1+2+3 and 1+3 clones (Figure S3 compare A and B), indicating an absence of central *GFP* reporter construct involvement in the formation of the early-replicating domain. A quantitative analysis revealed that the differences in RT between cell clones with one and two autonomous replicons were highly significant (*p*-value=0.01554 between (i) and (ii) and *p*-value=0.03571 between (ii) and (iii), Figure 5C). Overall, these results strongly suggest that two similar independent advanced replicons, 30 kb apart, can synergize to control the extent of the replication timing shift. We observed similar timing profiles 500 kb away from insertion site 1, in the 3’ proximal early-replicating domain (Figure 5D, compare with and +500 kb). Thus, the new timing of the early-replicating region extends to the 3’ proximal early domain.

**Figure 5:**
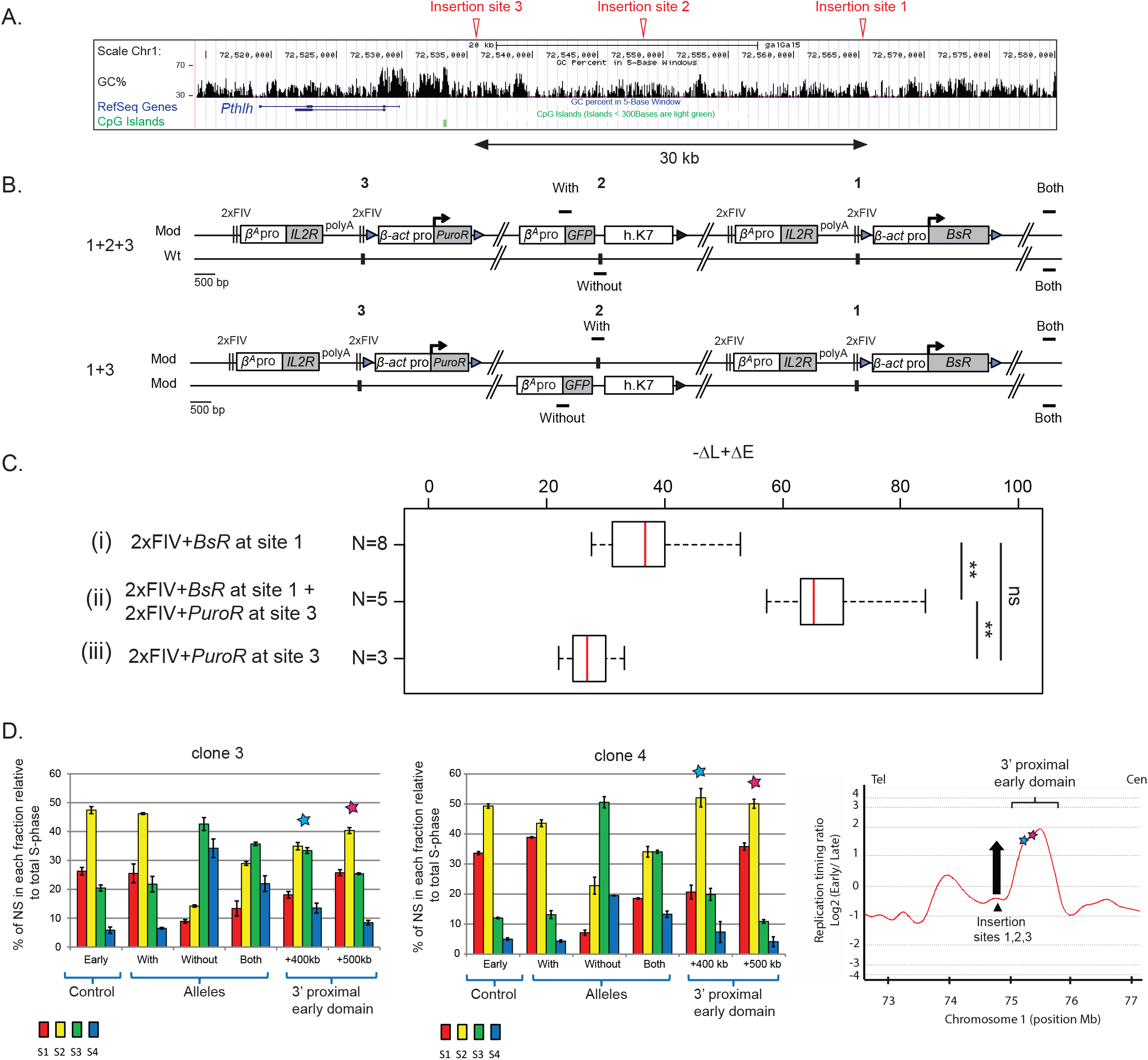
Two advanced replicons separated by 30 kb synergize to form a synthetic early replicated region. (A) UCSC genome browser visualization (Assembly Gallus_gallus-5.0/galGal5 Dec 2015) of a 65 kb window in the mid-late-replicated region of chromosome 1. Insertion sites 1, 2 and 3 are indicated with red arrows. Tracks showing CG percentage, annotated genes and CpG Islands are represented. (B) Cell lines containing two advanced replicons (2xFIV+*BsR* and 2xFIV+*PuroR*) inserted at sites 1 and 3, respectively, on the same chromosome and one *GFP* reporter construct composed of the *GFP* reporter gene under the control of the *β^A^-globin* promoter/origin linked to a 1.6 kb fragment of human chromosome 7 (h.K7) inserted at site 2, either on the same chromosome as 1 and 3 (1+2+3), or on the other chromosome (1+3). (C) Distribution of -ΔL+ΔE values for cell lines containing the 2xFIV+*BsR* transgene inserted at site 1 (i) or the 2xFIV+*PuroR* transgene inserted at site 3 (iii) and cell lines 1+3 and 1+2+3 (ii). (D) RT profiles of two 1+3 individual clones are shown (Figure S3B, clones 3 and 4). Quantifications with primer pairs located 400 kb and 500 kb downstream from site 1 in the 3’ proximal early-replicated domain (+400 kb, +500 kb). Error bars correspond to the standard deviation for qPCR duplicates. The RT profile along 4 Mb of chromosome 1 is shown on the right (genomic position: 72,650,000-76,840,000; 4.2 Mb; Assembly WUGSC 2.1/galGal3 May 2006). The three insertion sites (1, 2, 3) are indicated with a black triangle. The two new positions analyzed are indicated with a blue (+400 kb) and a pink (+500 kb) star.

### The formation of the early-replicating domain is associated with the spatial proximity of the two advanced replicons, which are 30 kb apart

Genome-scale studies have shown that RT is strongly correlated with chromatin interaction profile (Hi-C) in human cells, suggesting a spatial proximity of regions of similar RT within the nucleus (Ryba et al., 2010). This finding suggests that molecular interactions between genomic elements of similar RT are highly likely. We tested this hypothesis, by determining whether the synergy between the two advanced replicons leading to the formation of an early-replicating domain involved spatial proximity between the replicons concerned. Cre/*loxP* site-specific recombination has been successfully used in yeast and bacteria to assess the relative probabilities of different regions of the genome colliding with each another (Burgess and Kleckner, 1999; Hildebrandt and Cozzarelli, 1995). We therefore made use of this particular system to evaluate the spatial proximity between the two autonomous replicons located 30 kb apart. We assessed the capacity of the left recombination element located in the 2xFIV+*PuroR* construct at site 3 (loxP_LE, yellow triangle, Figure 6A) to recombine with the right recombination element in the 2xFIV+*BsR* construct at site 2 (loxP_RE, Figure 6A, green triangle). The percentage of chromosomes modified by this recombination (Figure 6B, large 1+3 excision) was quantified in cell clones with 1+2+3 and 1+3 configurations 1 h, 8 h, 24 h and 48 h after the induction of the Cre recombinase. We evaluated the frequency of random collisions, by constructing new clones in which the advanced replicon inserted at site 1 was replaced by a loxP_LE element alone (Figure 6A, 1(LoxP_RE)+3). We quantified the large excision product of the modified chromosome by semiquantitative PCR on the whole cell population with a specific primer pair (Figure 6B, For large excision and Rev large excision). The percentage of cells containing this product increased steadily over time, from 5 to 15% after 1 h to 45% to 89% of the cells after 48 h of Cre recombinase induction (Figure 6C and D). By contrast, only small amounts of this product were detected in three independent clones carrying the 1(LoxP_RE)+3 configuration, (from 4% after 1 h to 11% of the cells after 48 h of Cre recombinase induction; Figure 6C and D). Recombination rates varied by five to eightfold between the two types of configuration, as reported for yeasts harboring different combinations of *loxP* sites (Burgess and Kleckner, 1999). Overall, our data strongly suggest that molecular contact between the two constructs shifting RT increases, bringing the two *loxP* sites together. The formation of the early-replicating domain is, thus, linked to a spatial connection between the two advanced replicons located 30 kb apart, with the potential formation of a chromatin loop.

**Figure 6:**
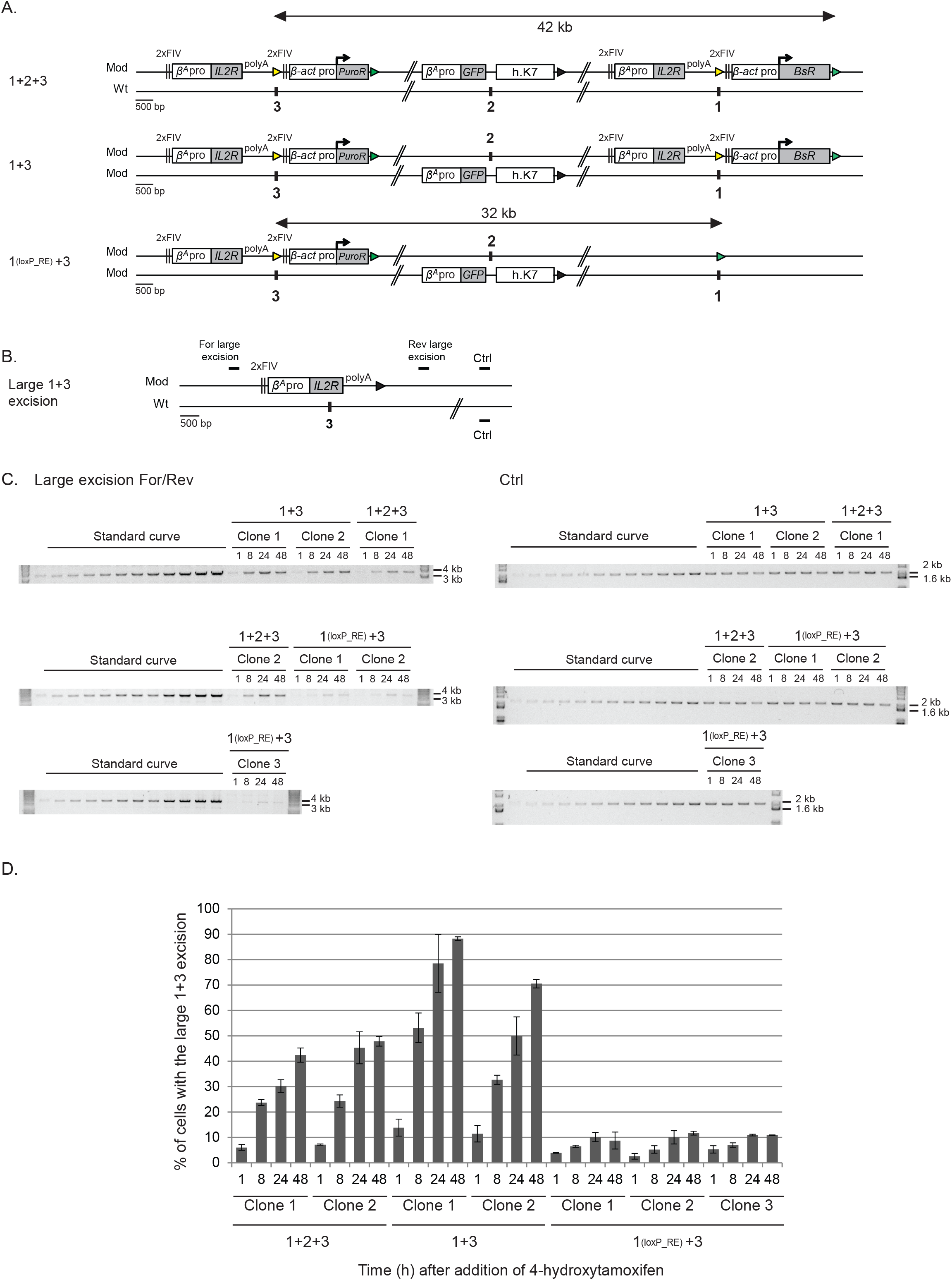
The formation of the early-replicating domain is associated with the spatial proximity of the two advanced replicons separated by 30 kb. (A) Clones 1+2+3 and 1+3 described in Figure 5B, together with control clones containing the 2xFIV+*PuroR* transgene inserted at site 3 and one loxP_RE sequence inserted at site 1 on the same chromosome and the *GFP* reporter construct inserted at site 2 on the other chromosome (1(LoxP_RE)+3) were tested for their capacity to recombine after induction of the Cre recombinase. Yellow and green triangles represent reactive loxP_LE and loxP_RE sites, respectively, and black triangles represent recombined inactive loxP sites. (B) Recombination between the upstream loxP_LE element (yellow triangle) inserted at site 3 and the downstream loxP_RE element (green triangle) inserted at site 1 leads to a large 1+3 excision product. (C) After 1, 8, 24 and 48 h of 4-hydroxytamoxifen treatment, genomic DNA was extracted and quantified by semi-quantitative PCR. PCR products specific for the large 1+3 excision (3.4 kb, in B thick black lines, For and Rev large excision, left gels) or used for normalization (1.9 kb, amplification from both chromosomes, in B thick black line, Ctrl, right gels) were run on a 0.8% or 1% w/v agarose gel, respectively, and stained with SYBR safe. The DNA size marker was a commercial 1 kb plus ladder. (D) The percentages of cells with the large 1+3 excision after 1, 8, 24 and 48 h of 4-hydroxytamoxifen treatment are shown for different cell lines.

### The close spatial proximity of the two advanced replicons is not associated with an increase in chromatin accessibility

We then investigated whether the synergy of the two autonomous replicons associated with their physical proximity affected the chromatin accessibility of these replicons or of one of the genomic regions located within the early-replicating domain. We addressed this question by quantifying, as previously described, nuclease-treated chromatin in 1+2+3 and 1+3 cell lines, by qPCR with specific primer sets specifically amplifying either one or both autonomous replicons (2xFIV+*BsR*, 2xFIV+*PuroR*), at the central *GFP* reporter transgene, at the three insertion sites or at control loci (Figure 7A). We compared chromatin accessibility in these cell lines with that in cell lines having only one 2xFIV+*BsR* construct (Figure 3C). The control regions behaved as previously described. However, we observed an increase in total nucleosome release at insertion site 3 relative to condensed chromatin controls, for all MNAse concentrations. The chromatin of this insertion site is, therefore, more accessible than that at sites 1 and 2 (Figure 7B, insertion site 3). The *BsR* module harbors the same pattern of accessibility as observed with one autonomous replicon. Nucleosomes embedded within the upstream 2xFIV element of the 2xFIV+*PuroR* construct (5’ 2xFIV-3) and within the *PuroR* gene were more susceptible to MNAse, regardless of MNAse concentration, than nucleosomes embedded at similar positions in the *BsR* module. The chromatin at this site is, therefore, also more exposed, probably reflecting the greater accessibility of the unmodified site 3. The *GFP* reporter construct in the middle of the early-replicating domain (1+2+3) or on the other, mid-late-replicating wt chromosome (1+3), released similar small amounts of material for all MNAse concentrations as were detected for heterochromatin regions. We can conclude from these results that two constructs composed of 5.6 kb of accessible chromatin and containing efficient replicators separated by 30 kb are sufficient to create an early-replicated domain when inserted into a naturally closed mid-late-replicating region. These two constructs can establish spatial connections without increasing the accessibility of their own chromatin, or of that located in the middle of the early-replicating domain.

**Figure 7:**
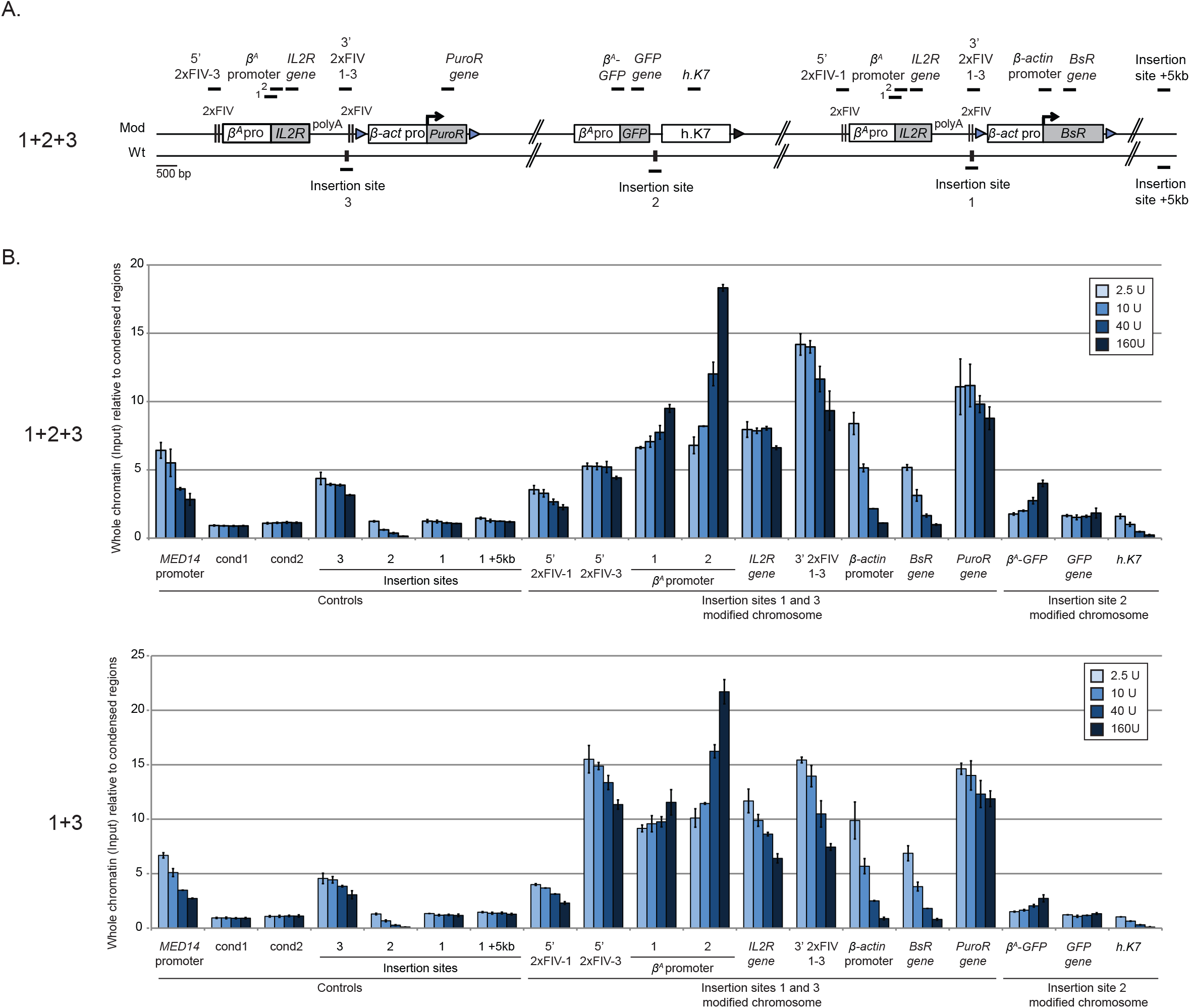
The close spatial proximity of the two advanced replicons is not associated with an increase in chromatin accessibility. (A) Clones 1+2+3 and 1+3 described in Figure 5B were used to assess chromatin accessibility within the transgenes. The positions of the amplicons used for quantification in each shifting construct individually (thick black lines; 5’ 2xFIV-1 or −3, *β-actin* promoter, *BsR* gene, *PuroR* gene), in both shifting constructs (*β^A^* promoter 1 and 2, *IL2R* gene, 3’ 2xFIV1-3) or in the *GFP* reporter construct (*β^A^*-*GFP*, *GFP* gene, h.K7) are shown. The insertion sites on the wt chromosome (insertion sites 1, 2 and 3) and a genomic region located 5 kb downstream from the insertion site (insertion site +5kb) were used as controls for the targeted genomic region. (B) Quantifications by real-time qPCR of total chromatin (input) extracted from two clones of 1+2+3 and 1+3 cell lines, after digestion with micrococcal nuclease (MNAse, 2.5 U, 10 U, 40 U, 160 U/mL). Error bars indicate the standard deviation for qPCR triplicates, for two independent clones. Data are presented as total chromatin input relative to the condensed genomic regions (cond1 and cond2) used for the normalization.

## Discussion

The DNA replication timing program has emerged as a system that integrates genome regulation and the three-dimensional organization of the genome. These observations underlie a model (the replication domain model) in which each RT domain acts as a regulatory unit, determining when the replicons within its boundaries can fire. We tested this model, by using constructs containing critical *cis*-elements known to regulate origin firing to disturb mid-late- and late-replicating domains in a specific manner. Based on the work described here, we identified four important principles underlying the establishment and maintenance of domains of early and late replication: (1) A strong origin associated with specific chromatin features can significantly advance the RT of a mid-late-replicating region; (2) Two replicators carrying timing information can act in cooperation if located in close proximity; (3) The replication timing of late-replicating domains can be advanced locally by a strong autonomous replicon, but the final RT is also influenced by chromosomal context; (4) Strong autonomous replicons can act in synergy even when separated by 30 kb, and they tend to come into close spatial proximity.

### Molecular mechanisms shaping efficient replicons carrying timing information

Studies in *S. cerevisiae* have suggested that the correct temporal activation of origins during S-phase involves the activation of early-firing origins, because these origins have a greater “accessibility” to firing factors, which are present in limiting amounts. After firing, the limiting factors are released and can bind to and activate less accessible origins, and so on (Douglas and Diffley, 2012). This theory suggests that the RT of a region directly reflects the accessibility to these limiting factors of specific regions bound by pre-RCs. In vertebrate cells, there is a strong correlation between early replication and the accumulation of open chromatin marks, consistent with a role for the openness of chromatin in the recruitment of these potential limiting factors (Picard et al., 2014; Ryba et al., 2010). We previously identified a combination of *cis*-elements constituting an independent replicon with the capacity to advance RT significantly for a portion of a mid-late-replicating region (Hassan-Zadeh et al., 2012). This construct, although synthetic, contains only promoters and regulatory motifs present in the chicken genome (one tissue-specific promoter, one constitutive promoter and four FIVs), linked to either the *IL2R* reporter gene or the *blasticidin* gene selection cassette. Moreover, autonomous replicon activity is driven only by endogenously expressed *trans*-factors. We first investigated in more detail the chromatin structure of each independent unit. We identified a number of different features, highlighting the potential diversity of *cis*-elements involved in RT control in vertebrates. One of the elements identified contains the *β^A^-globin* erythroid tissue-specific promoter which cannot drive transcription in the DT40 lymphoid cell line. Accordingly, this promoter does not carry the histone marks typical of active promoters (H3K4me3). However, we nevertheless observed an enrichment in H3K4me2 at the promoter and USF-flanking binding sites, together with greater chromatin accessibility at USF sites. The other element contains the constitutive *β-actin* promoter linked to a gene for selection. This strong origin/promoter facilitates the local opening of chromatin structure and the acquisition of H3K4me3 and H3K9K27ac marks. These findings are consistent with the recent observation that the synthetic activation of transcription, but not nuclear repositioning, shifts the RT of several mid-late-replicating loci to mid-early in mouse ES cells (Therizols et al., 2014). Our first observation shows that, at least at this mid-late-replicated chromosomal position, origins located within the *β^A^-globin* and *β-actin* promoters are highly efficient, because they were able to advance RT locally in a population-based timing assay. We were unable to estimate the fraction of cells in which these two replicators were activated, but the significant shift in RT toward earlier replication strongly suggests that they are activated in a large proportion of cells. Based on genome-wide studies and our recent genetic study, we suggest that most vertebrate origins consist of a G-quadruplex motif linked to an as yet undefined *cis*-element, and that replicators overlapping a promoter tend to be the most effective (Besnard et al., 2012; Cayrou et al., 2012; Valton et al., 2014). Both the *β^A^-globin* and *β-actin* promoters/origins contain a G-quadruplex motif and locally induce SNS enrichment when inserted ectopically, thereby satisfying the criteria for a “canonical strong origin” (Hassan-Zadeh et al., 2012).

Statistical analyses showed that these origins together with *cis*-elements controlling chromatin structure could act in cooperation when located close together, by providing more effective timing information. This result supports the hypothesis that early-replicating domains result from the clustering of origins associated with the opening of the chromatin structure. Consistent with this hypothesis, genome-wide mapping of replication origins by the SNS method in human and mouse cells has shown the clustering of efficient replication origins within early-replicated domains, forming small initiation zones of several kilobases, mostly located around TSS containing a CpG island (CGI) (Cayrou et al., 2015; Picard et al., 2014). These endogenous structures are reminiscent of the organization of our two linked RT shift-inducing constructs.

### Early replicators can advance replication timing locally in late-replicating domains

We investigated the possible dominance of late-replicating domains over replicators carrying early timing information, by assessing the capacity of the same dissected combination of *cis*-elements to function as an independent replicon in genomic regions that are naturally replicated in late S-phase. This construct had similar impacts on RT at three chromosomal loci (one replicated in mid-late S-phase and two in late S-phase), demonstrating the robustness of the signal embedded in this specific construct. With respect to the replication domain model, these findings suggest that any type of chromosomal environment is permissive for the formation of an autonomous advanced replicon. However, the observation that the final RT for the modified region was more advanced for the region normally replicated in mid-late S-phase than for that normally replicated in late S-phase suggests that there may be two layers of regulation, at the local and large scales. Both these levels of regulation must be taken into account if we are to understand how large replication domains are constructed. Our data suggest that late-replicating domains are also defined by a lack of signals associated with autonomous early replicons. Consistent with this hypothesis, one mechanism proposed for the regulation of late-replicated regions associated with common fragile sites involves a low density of replication initiation events (Letessier et al., 2011). Less abundant but nevertheless efficient origins can also be detected in late-replicating domains by the SNS method. A recent study investigated the role of H4K20me marks in controlling late-firing origins. It showed that the conversion of K20me1 to K20me3 enhances ORCA recruitment and MCM loading at already defined origins required for the correct replication of heterochromatin (Brustel et al., 2017). Interestingly, the loss of Su4-20h, which is responsible for this methylation, delays RT for 15% of the mouse genome. Late origins are, therefore, also defined by a specific chromatin organization facilitating the coordination of late firing and counteracting the overall repressive effect of heterochromatic regions without providing early-firing signals.

### Robust early-replicated domains can be formed by synergy between efficient early replicators separated by several tens of kilobases

We investigated the formation of early-replicating domains further, by inserting two identical strong combinations of early origins into the same chromosome, at sites 30 kb apart. This distance is within the range of average spacings between strong initiation sites or zones detected by the SNS method in early-replicated domains. To our surprise, we found that these two inserted constructs advanced RT to a much greater extent when inserted together than when inserted alone. We found that the presence of the two constructs favored physical contact between these two remote regions. We suspect that this spatial connection is required for the synergic effect on RT, although we have no definitive proof that this is the case. Interactions between active promoters have been observed by several methods, including the recent production of an ultrahigh-resolution Hi-C map during neural differentiation in mice (Bonev et al., 2017; Li et al., 2012). These results are consistent with our observations, because our construct shown to act synergically contains the constitutive *β-actin* promoter. In *S. cerevisiae* the Fkh1/2 transcription factors have been shown to be directly involved in the early firing of a large group of origins (Knott et al., 2012).

It has recently been shown that this function involves direct interaction with Dbf4, the regulatory unit of the essential DDK firing factor (Fang et al., 2017). Based on the dimerization capacity of Fkh1/2 and the observation that early firing origins associated with Fkh1/2 tend to cluster, it was suggested that this clustering might even increase the efficiency of Fkh1/2 for the local recruitment of Dbf4 to early replication factories (Knott et al., 2012). We suggest that the close proximity of two strong autonomous replicators carrying timing information here locally increased the efficiency of limiting firing factor recruitment in a similar manner. Most efficient early-firing origins are associated with active promoters containing CGI. This model can, therefore, be adapted to genome-wide observations, validating our synthetic approach.

## Acknowledgments

The authors thank the members of the laboratory of M-N.P for useful insights and discussions. We thank G.Felsenfeld and R.Veitia for critical reading of the manuscript. We thank J-F Bercher for his help in defining the most accurate quantitative method for measuring RT shifts. We thank Griselda Wentzinger for performing cell sorting at ImagoSeine Institut Jacques Monod platform. This work was supported by grants from the Association pour la Recherche sur le Cancer (Equipe Labellisée) and the Agence Nationale pour la Recherche (ANR-15-CE12-0004-01). M-N.P. is supported by Inserm.

## Author Contributions

Conceptualization: CB, BD and M-NP; Methodology: CB and M-NP; Validation: CB, BD and M-NP; Investigation: CB, SC, AC, ML, CG; Writing-original Draft: CB and M-NP; Visualization: CB; Supervision: CB, BD and M-NP; Project administration: M-NP; Funding Acquisition: M-NP.

## Declaration of Interests

The authors declare no competing interests.

## STAR Methods

### Plasmid construction

The targeting vectors for homologous recombination in DT40 cells were constructed with the multisite Gateway Pro kit (Thermo Fischer Scientific), as previously described (Hassan-Zadeh et al., 2012). The same entry vectors as before were used for the 5′ and 3′ target arms for specific insertion at site 1 in the mid-late-replicating region (chr1:74,813,240 Assembly WUGSC 2.1/galGal3 May 2006, chr1:72,565,520 Assembly Gallus_gallus-5.0/galGal5 Dec 2015) (Hassan-Zadeh et al., 2012). New arms were prepared for specific targeting at sites 2 (chr1: 74,795,973 Assembly WUGSC 2.1/galGal3 May 2006, chr1: 72,548,589 Assembly Gallus_gallus-5.0/galGal5 Dec 2015) and 3 (chr1: 74,783,147 Assembly WUGSC 2.1/galGal3 May 2006, chr1: 72,536,060; Assembly Gallus_gallus-5.0/galGal5 Dec 2015) in the mid-late-replicating region, and at sites 1 (Late1 chr1:72,616,319 Assembly WUGSC 2.1/galGal3 May 2006, chr1:70,523,649 Gallus_gallus-5.0/galGal5 Dec 2015) and 2 (Late2 chr1:182,214,693 Assembly WUGSC 2.1/galGal3 May 2006, chr1:177,936,192 Gallus_gallus-5.0/galGal5 Dec 2015) in the late regions. The 5’ and 3’ target arms for homologous recombination were amplified from DT40 genomic DNA with the following primer pairs: 5’arm for_ML2: CCAAACCAGGCCACTCTTAGT; 5’arm rev_ML2: AGTCACTTGGCATAAATAAGAAGCC and 3’arm for_ML2: CTGAGCAGGAAGGGAAACGA; 3’arm rev_ML2: CCATAGTGCAGACCTGGCAT, to yield 2,163 bp and 2,068 bp amplicons, respectively, for integration at mid-late site 2; 5’arm for_ML3: ACACACTCACCTCCTGCCTT; 5’arm rev_ML3: AGATCTCAGTCCTGCCAGCA and 3’arm for_ML3: AAGTTCGTAATACACAACCTTGAC; 3’arm rev_ML3: GCTTCCCCGCTTCTTCCCTA, to yield 2,054 bp and 2,062 bp amplicons, respectively, for integration at mid-late site 3; 5’arm for_L1: CAGTGAAACACAGGAGGAACA; 5’arm rev_L1: TAACTCCAAGAACGATCACTGC and 3’arm for_L1: GGAAATGTCTTGAATCTCACAAGG; 3’ arm rev_L1: ATGCCCACCAGTGTCCATAA, to yield 2,244 bp and 2,137 bp amplicons, respectively, for integration at late site 1; 5’ arm for_L2: ACTTGTTGAGCCTTTATGGAGAAC; 5’ arm rev_L2: CGGTGTTACAGAGGAGTAAACTGA and 3’ arm for_L2: ACCTTATGCATTTCGTTCCTATGT; 3’ arm rev_L2: TAAGAAGAGAGATGGGGATCAAAC, to yield 1996 bp and 2001 bp amplicons, respectively, for integration at late site 2.

We used four entry vectors to generate the new 2xFIV+*BsR* construct inserted at site 1: two entry vectors containing the 5′ and 3′ target arms for specific insertion at site 1 and two new entry vectors. The first contained two copies of FIV linked to the *β^A^* promoter (2x FIV_UP). The 2XFIV entry vector was used as a template for PCR amplification with the following primers: 2XFIV-for: 5′GGGGACAACTTTGTATACAAAAGTTGAGGTGGCACGGGATCGCTTTCCTAGGTGGCACGGGATCGCTTTCCTCTGCCCACACCCTCCTG3′; Rev_SpeKpn: 5′GGGGACAACTTTGTATAGAAAAGTTGGGTGGGTACCACTAGTGATGATCCGTCATCCAGAC ATG3′. The second entry vector (2xFIV_*BsR*) contained the 2xFIV elements upstream from the *blasticidin* resistance (*BsR*) gene cassette, flanked by *loxP* sites. This construct was made by amplifying the *β-actin* promoter of the *BsR* gene cassette from the pLox*BsR* vector (Arakawa et al., 2001) such that *Xho*I and 2xFIV sequences were added upstream from the promoter with the following primers: 2XFIV_*BsR_*for: 5′GCTCGAGAGGTGGCACGGGATCGCTTTCCTAGGTGGCACGGGATCGCTTTCCTGTGAGC

CCCACGTTCTGCTT3′. *BsR*_rev: 5′CTTCTCTGTCGCTACTTCTAC3′. The PCR product was inserted into the *BsR* entry vector, between the *Xho*I site located upstream from the *β-actin* promoter and the *Bgl*II site at the end of the promoter. The PCR product was introduced into the corresponding entry vector by ligation with T4 DNA ligase (BioLabs), and the 2xFIV_*BsR* entry vector was selected after sequence verification. The corresponding final vector was generated by recombining compatible *att* sites between the entry vectors, with LR clonase. For electroporation, the final vector was linearized with *Sca*I.

We used four entry vectors to generate the 2xFIV+*PuroR* construct inserted at site 3. We used two entry vectors containing the 5′ and 3′ target arms for specific insertion at site 3: the previously described 2x FIV_UP entry vector and a new entry vector (2xFIV_*PuroR*) containing the 2xFIV sequence upstream from the *puromycin* resistance (*PuroR*) gene cassette, flanked by *loxP* sites. This construct was made by amplifying part of the entry vector from the previously described 2xFIV_*BsR* vector such that the *attB* sites were flanked by the *Xho*I restriction site and 2xFIV sequences on one side of the recombined region and the *Xho*I sequence on the other, with the following primer pair: XhoI and 2x FIV_entry vector for: 5’GACTCTCGAGAGGAAAGCGATCCCGTGCCACCTAGGAAAGCGATCCCGTGCCACCTGCTAGCCCTGATCAATAACT3’ and XhoI entry vector rev: 5’AAGTCCTCGAGCGTATTACAATTCACTGGCCGT3’. The *β-actin* promoter associated with the *PuroR* gene cassette was produced by *Xho*I digestion of the pLoxPuro plasmid (Arakawa et al., 2001). The final 2xFIV_*PuroR* entry vector was generated by *Xho*I digestion of the PCR product and ligation with T4 DNA ligase. The corresponding entry vector was selected after sequence verification. For electroporation, the final vector was linearized with *Pvu*I.

We used four entry vectors to generate the *GFP* reporter construct inserted at site 2. We used two entry vectors containing the 5′ and 3′ target arms for specific insertion at site 2, one entry vector previously described and containing a similar *BsR* selection cassette flanked by two *loxP* sites (Hassan-Zadeh et al., 2012) and one new entry vector *(β^A^-GFP*-h.K7). This entry vector contained the *β^A^-globin* promoter sequence amplified from the previously described 2x FIV_UP entry vector with the following primer pair: *β*^A^_for: 5’CTGCCCACACCCTCCTG3’ and *β*^A^_rev: 5’ TTCCTGACCCTTGGGACCA3’, the *GFP* gene sequence amplified from the peGFP plasmid (clontech 6085-1) with the following primer pair: *GFP*_for: 5’GGGTCAGGAAATGGTGAGCAAGGGCGAGG3’ and *GFP*_rev: 5’TGAAGCAGCATTTACGCCTTAAG3’ and part of human chromosome 7 (chr7:26,873,165-26,874,805; Assembly (GRCh38/hg38)Human Dec. 2013) amplified from human genomic DNA with the following primer pair: h.K7_for: 5’TGCTGCTTCATTTCTGCTCTC3’ and h.K7_rev: 5’GCAGAGCCAGAGTCCAAGAG3’. The *β*^A^_for and h.K7_rev primers were associated with the corresponding *attB* sites for Gateway recombination. These three DNA fragments were combined with the In-Fusion HD cloning plus kit (Clontech), introduced into the corresponding entry vector with BP clonase (Thermo Fisher Scientific) and selected after sequence verification. For electroporation, the final vector was linearized with *Pvu*I.

The *loxP* RE sequence was inserted into site 1 by the insertion of a larger construct composed of the *loxP* RE-h.K7+*BsR* elements followed by site-specific Cre recombinase excision. This construct was generated by the association of two entry vectors containing the 5′ and 3′ target arms for specific insertion at site 1, a previously described entry vector containing a similar *BsR* selection cassette (Hassan-Zadeh et al., 2012) and a new entry vector (loxP RE-h.K7). This entry vector contained part of the human chromosome 7 sequence amplified from human genomic DNA with the following primer pair containing the upstream *loxP* RE sequence: loxP RE-h.K7_for 5’GGATCCATAACTTCGTATAGCATACATTATACGAACGGTAACTTGAGCCCAGGAGTTCGA3’ and h.K7_rev 5’ AGAGTTCCAACCCCAGCCTC3’. The loxP RE-h.K7_for and the h.K7_rev primers were associated with the corresponding *attB* sites. The PCR products were introduced into the corresponding entry vector with BP clonase (Thermo Fisher Scientific) and selected after sequence verification. For electroporation, the final vector was linearized with *Pvu*II.

### Cell culture and transfection

DT40 cells were grown in RPMI 1640 Glutamax (Thermo Fisher Scientific) containing 10% FBS, 1% chicken serum, 0.1 mM β-mercaptoethanol, 200 U/mL penicillin, 200 µg/mL streptomycin and 1.75 µg/mL amphotericin B, at 37°C, under an atmosphere containing 5% CO_2_. We transfected DT40 cells as previously described (Hassan-Zadeh et al., 2012). Cell clones were selected on media containing a final concentration of 20 µg/ml blasticidin or 1 µg/ml puromycin, depending on the resistance gene carried by the transgene. Genomic DNA was extracted from cells in lysis buffer (10 mM Tris pH8.0; 25 mM NaCl; 1 mM EDTA and 200 µg/mL proteinase K). Clones into which the plasmid DNA was integrated were screened by PCR with primer pairs designed to bind on one side of the insertion site such that one primer bound within the construct and the other primer bound just upstream or downstream from the arm used for recombination, as previously described (Hassan-Zadeh et al., 2012)(Figure S5 and Table S3). The insertion of constructs into the same chromosome or the other chromosome was determined by long-range PCR (Figure S6). For each clone tested, genomic DNA was extracted with the DNeasy Blood & Tissue kit (Qiagen). We designed primer pairs amplifying in the part of the genome between constructs to control for DNA extraction quality (Ctrl), and primer pairs amplifying the two tested constructs to test the insertions (Table S3). The LR 1+2 primer pair was used to amplify the h.K7 part of the *GFP* reporter construct inserted at site 2 and the *IL2R* gene within the 2xFIV+*BsR* construct inserted at site 1. The LR 2+3 primer pair was used to amplify the *PuroR* gene within the 2xFIV+*PuroR* construct inserted at site 3 and the *GFP* gene within the *GFP* reporter construct inserted at site 2. The LR 1+3 primer pair was used to amplify the *PuroR* gene within the 2xFIV+*PuroR* construct inserted at site 3 and the *IL2R* gene within the 2xFIV+*BsR* construct inserted at site 1, regardless of the presence or absence of the central construct. The LR 1(loxP_RE)+3 primer pair was used to amplify the *PuroR* gene within the 2xFIV+*PuroR* construct inserted at site 3 and the loxP_RE element inserted at site 1, regardless of the presence or absence of the central construct. PCR was performed with the Long PCR Enzyme Mix (Thermo Fisher Scientific) under following conditions: initial denaturation at 94°C for 3 minutes, followed by 10 cycles of 95°C for 20 s, 68°C for 14 minutes, followed by 20 cycles of 95°C for 20 s, 68°C for 18 minutes and a final extension phase of 10 minutes at 68°C. Quantitative PCR (qPCR) was carried out for each clone, on 4 ng of genomic DNA, with a primer binding to the transgene (*β*^A^-*GFP* on the *GFP* reporter construct, With on 2xFIV+*BsR* or 2xFIV+*PuroR*, With on *BsR* and LoxP_RE on the loxP site inserted at site 1), and a primer binding close to insertion site 1 and amplifying both alleles (Both or insertion site 1 + 5 kb), to confirm that only one copy of the transgene had been inserted (Table S2 and S3).

### Cre-loxP excision

In this study, we used the DT40 Cre1 subclone, which constitutively expresses a tightly regulated Cre recombinase fused to a mutated estrogen receptor (Mer) (Arakawa et al., 2001). This inactive Mer-Cre-Mer fusion protein can be transiently activated in the presence of 4-hydroxytamoxifen, resulting in the efficient excision of genomic regions flanked by two recombination signals (*loxP* sites) inserted in the same direction. We also used a modified inducible Cre recombination system involving two different mutant loxP sites (loxP_RE and loxP_LE). After Cre recombination, these two mutants are converted into a new nonfunctional loxP site (loxP_RE+LE) that is not recognized by the Cre recombinase, thereby preventing additional recombination events (Arakawa et al., 2001). For the excision of the genomic DNA flanked by loxP sites, we treated 3 x 10^5^ cells with 5 μM 4-hydroxytamoxifen (Sigma Aldrich) for two days. Subclones were obtained by plating dilutions of the treated cell suspension at a density of 50, 150, and 1500 viable cells per 10 ml in 96-well flat-bottomed microtiter plates. Genomic DNA was extracted from single subclones and analyzed by PCR with specific primer pairs (Table S3). We assessed the excision of the *BsR* selection cassette in clones with the *GFP* reporter construct inserted at site 2, with a primer pair amplifying the polyA sequence of the both *BsR* and *GFP* genes and the downstream part of the 3’ arm of insertion site 2 (Figure S5D, primer pair #3-4). All clones were cultured for 72 h in selective media containing the appropriate antibiotic, to confirm PCR results.

For the large 1+3 excision kinetic, 3 x 10^5^ cells were treated with 5 μM 4-hydroxytamoxifen for 1, 8, 24, 48 h before genomic DNA extraction with the DNeasy Blood & Tissue kit (Qiagen). The large 1+3 excision was quantified by semiquantitative PCR with the large excision primer pair and the Ctrl primer pair used for normalization (Figure 6B and Table S3). Calibration was based on a standard curve generated with various amounts of genomic DNA (16 ng to 160 ng) obtained from a previously isolated clone in which the large recombination had occurred. For each set of excision kinetic conditions, 64 ng or 160 ng of genomic DNA was tested (for amplification with the Ctrl or the large excision primer pair, respectively). PCR was performed with the Herculase II fusion DNA polymerase (Agilent Technologies) under the following conditions: initial denaturation at 98°C for 4 minutes, followed by 25 cycles of 98°C for 30 s, 57°C for 45 s, 72°C for 2 minutes, and a final extension phase for 3 minutes at 72°C. After migration of the PCR products on a 0.8% agarose gel and SYBR Safe staining, the mean intensity of each signal was quantified with Image J, corrected with the local background mean intensity and normalized with the mean intensity of the signal obtained in the control amplification.

### Replication timing analysis

The RT experiments presented in Figures 4, 5 and S1 to S3 were performed as previously described (Hassan-Zadeh et al., 2012). Briefly, about 10^7^ exponentially growing cells were pulse-labeled with BrdU for 1 h and sorted into four S-phase fractions, from early to late S phase. The collected cells were treated with lysis buffer (50 mM Tris pH 8.0; 10 mM EDTA pH 8.0; 300 mM NaCl; 0.5% SDS, 0.2 mg/ml of freshly added proteinase K and 0.5 mg/ml of freshly added RNase A), incubated at 65°C for 2 h and stored at −20°C, in the dark. Genomic DNA was isolated from each sample by phenol-chloroform extraction and alcohol precipitation and sonicated four times for 30s each, at 30s intervals, in the high mode at 4°C in a Bioruptor water bath sonicator (Diagenode), to obtain fragments of 500 to 1000 bp in size. The sonicated DNA was denatured by incubation at 95°C for 5 minutes. We added monoclonal anti-BrdU antibody (BD Biosciences) at a final concentration of 3.6 μg/ml in 1x IP buffer (10 mM Tris pH 8.0, 1 mM EDTA pH 8.0, 150 mM NaCl, 0.5% Triton X-100) containing 7 mM NaOH. We used 30 μl or 50 µl of protein-G-coated magnetic beads (from Ademtech or Thermo Fisher Scientific, respectively) per sample to pull down the anti-BrdU antibody. Beads and BrdU-labeled nascent DNA were incubated for 2-3 hours at 4°C, on a rotating wheel. The beads were then washed once with 1x IP buffer, twice with wash buffer (20 mM M Tris pH 8.0, 2 mM EDTA pH 8.0, 250 mM NaCl, 0.25% Triton X-100) and then twice with 1x TE buffer pH 8.0. The DNA was eluted by incubating the beads at 37°C for 2 h in 250 µl 1x TE buffer pH 8.0, to which we added 1% SDS and 0.5 mg/ml proteinase K. DNA was purified by phenol-chloroform extraction and alcohol precipitation and resuspended in 50 µl TE.

Genome-wide RT analyses were performed as previously described (Hassan-Zadeh et al., 2012) except that S-phase was divided into two fractions. For DNA microarrays, immunoprecipitated nascent strands were amplified by whole-genome amplification (WGA; Sigma) to obtain sufficient DNA. After amplification, early and late nascent strands were labeled with Cy3 and Cy5 ULS molecules (ULS array CGH labeling kit; Kreatech) as recommended by the manufacturer. Hybridization was performed according to the manufacturer’s instructions, on 46180K Chicken microarrays (Chicken Genome CGH Microarray 46180K, custom microarray design, genome reference Gallus gallus V3 May 2006) covering the whole chicken genome, with one probe every 5.6 kb. Microarrays were scanned with an Agilent High-Resolution C Scanner, using a resolution of 3 µm and the autofocus option. Features were extracted with Feature Extraction 11.5.1.1 software (Agilent Technologies), with the CGH_107_Sep09 protocol. For each experiment, the raw datasets were automatically normalized by the Feature extraction software. Agilent Genomic Workbench 6.5 software was used for analysis. The log_2_-ratio timing profiles were smoothed with the moving average of the Agilent Genomic Workbench 6.5 software, with the triangular algorithm and 200 kb windows. The profile shown was obtained by combining intra- and interarray replicates with the error model algorithm for four hybridization experiments.

### Flow cytometry analysis

After BrdU incorporation, DT40 cells were washed twice with PBS, fixed in 75% ethanol and stored at - 20°C. On the day of sorting, fixed cells were resuspended at a final concentration of 2.5 x 10^6^ cells/mL in 0.1% IGEPAL in PBS (Sigma, CA-630), 50 μg/ml propidium iodide and 0.5 mg/ml RNase A, and incubated for 30 minutes at room temperature. Cells were sorted with an INFLUX 500 cell sorter (Cytopeia, BD Biosciences). Four fractions of S-phase cells (S1-S4), each containing 5×10^4^ cells, were collected and further treated for locus-specific RT analyses. Two fractions of S-phase cells (S1-S2), each containing 10^5^ cells were collected and further treated for genome-wide RT analyses.

### MNAse digestions

We cross-linked 30×10^6^ exponentially growing cells by incubation for 5 min with 1% (v/v) freshly prepared formaldehyde (Thermo Fisher Scientific) at room temperature. Fixation was stopped by adding glycine to a final concentration of 0.125 M and incubating for five minutes at room temperature, with stirring. Cells were then washed three times in cold PBS and their nuclei were extracted in 3 mL of lysis buffer (10 mM Tris-HCl pH 7.5, 10 mM NaCl, 3 mM MgCl_2_, 0.2% Triton X-100, 0.5 mM EGTA pH 8.0, 1 mM DTT, 1x protease inhibitor cocktail (Sigma, P8340)) for 5 minutes at 4°C, centrifuged and resuspended in digestion buffer (10 mM Tris-HCl pH 7.5, 10 mM NaCl, 3 mM MgCl_2_, 1 mM CaCl_2_, 1x protease inhibitor cocktail (Sigma, P8340)). Micrococcal Nuclease (MNase; Thermo Fisher Scientific, EN0181) digestions were performed for 15 minutes at 37°C, using either a final concentration of 10 U/mL for ChIp analyses or a series of four exponentially increasing concentrations of MNase (2.5, 10, 40 and 160 U/mL) for chromatin accessibility analyses. The reaction was stopped by adding 0.1 volume of stop buffer (200 mM EDTA pH 8.0, 40 mM EGTA pH 8.0). For ChIp analyses, MNase digestion conditions were established so as to produce mostly mono-to hexanucleosome fragments of 150 bp to 1000 bp.

Chromatin was then diluted by a factor of two in 2x complement buffer (40 mM Tris-HCl pH 8.0, 300 mM NaCl, 2% Triton X-100) and sonicated 20 times for 30s each, at 30s intervals in the high mode at 4°C in a Bioruptor water bath sonicator (Diagenode). For chromatin accessibility analyses, the lowest MNAse concentration generated a mixture of oligo-, di- and mononucleosomes, whereas the highest concentration produced mostly mononucleosomes (Figure S4A and B, respectively). After digestion, DNA molecules were extracted in phenol-chloroform and precipitated with ethanol and were then subjected to a size selection process with SPRIselect beads (Beckman Coulter) using a 0.5X ratio to remove DNA molecules of more than 1000 bp in length (Figure S4). The MNAse titration approach can be used to determine whether nucleosome release requires low or high levels of MNAse. Low levels of MNAse release large numbers of nucleosomes in accessible regions, leading to higher levels of DNA molecules. Nucleosomes embedded in less accessible regions are less likely to be released with such low levels of MNAse. Moreover, this approach provides additional information about the accessibility of loci. Indeed, highly inaccessible or condensed regions are not sensitive to higher concentrations of MNAse, whereas the chromatin of accessible regions is rapidly digested under the same conditions.

### Chromatin immunoprecipitation

Immunoprecipitation was performed overnight at 4°C, in a final volume of 200 µl of 1x IP buffer (20 mM Tris-HCl pH 8.0, 2 mM EDTA pH 8.0, 150 mM NaCl, 1% Triton-X100 and 0.1% SDS) on an amount of chromatin corresponding to 10 μg of DNA, with anti-trimethylated K4H3 (AbCam, ab8580), anti-dimethylated K4H3 (Millipore # 07-030) or anti-acetylated K9K27H3 (Millipore # 06-599) antibodies, according to the manufacturer’s recommendations. Immunocomplexes were pulled down with 50 µl of protein-G-coated magnetic beads (Thermo Fisher Scientific, Dynabeads Protein G) per sample. Beads and immunocomplexes were incubated for two hours at 4°C, on a rotating wheel. The beads were then washed once with 1x IP buffer, twice with wash B buffer (20 mM M Tris pH 8.0, 2 mM EDTA pH 8.0, 250 mM NaCl, 0.25% Triton X-100) and then twice with 1x TE buffer pH 8.0. For the anti-trimethylated K4H3 antibody, the wash buffer contained 500 mM NaCl, and an additional washing step was performed with buffer C (10 mM Tris-HCl pH 8.0, 1 mM EDTA pH 8.0, 1% sodium deoxycholate, 1% NP40, 250 mM LiCl). The DNA was eluted by incubating the beads for 2 h at 37°C with 250 µl 1x TE buffer pH 8.0, to which we added 1% SDS and 0.5 mg/ml proteinase K. Cross-linking was reversed by overnight incubation at 65°C and samples were further treated with 10 µg of RNase A for 15 minutes at 37°C, and with 20 µg of proteinase K for 1 h at 56°C. DNA was purified by phenol-chloroform extraction, precipitated in alcohol and resuspended in 100 µl TE.

### RNA extraction and reverse transcription

Total RNA were extracted from 5×10^6^ cells with the Nucleospin RNA kit (Macherey Nagel). The integrity of the extracted RNA was assessed with an Agilent 2100 bioanalyzer, and 20 µg of total RNA was then treated with 4 units of DNAse I (BioLabs) for 1 h at 37°C. The enzyme was inactivated by adding 5 mM EDTA and incubating the reaction mixture for 10 min at 75°C. The RNA was then purified by phenol-chloroform extraction and ethanol precipitation. Reverse transcription reactions (RT-PCR+) were then performed with 5 µg of RNA and random hexamers (BioLabs), using the Superscript III Reverse Transcriptase (Thermo Fisher Scientific) according to the manufacturer’s instructions. Negative controls (RT-PCR-) were performed with the same procedure, but without the addition of reverse transcriptase. The comparison of RT-PCR+ and RT-PCR-samples was used to validate DNAseI treatment and the complete digestion of the genomic DNA in the RNA samples.

### Real-time PCR quantification of DNA

The Roche Light Cycler 2.0 detection system and the Absolute Q-PCR SYBR green capillary mix (Thermo Fisher Scientific) were used for the real-time PCR quantification of BrdU-labeled newly synthesized strands (NS) or genomic DNA extracted from 4-hydroxytamoxifen-treated clonal cell lines. For all reactions, real-time PCR was performed under the following cycling conditions: initial denaturation at 95°C for 15 minutes, followed by 50 cycles of 95°C for 15 s, 60°C for 30 s, 72°C for 20 s, and fluorescence measurement. Following PCR, a thermal melting profile was used for amplicon identification.

The Roche Light Cycler 480 detection system and the LightCycler 480 SYB Green I Master mix (Roche Applied Science) were used for the real-time PCR quantification of ChIp DNA, MNAse-digested DNA and cDNA. For all reactions, real-time PCR was performed under the following cycling conditions: initial denaturation at 95°C for 5 minutes, followed by 50 cycles of 95°C for 10 s, 61°C for 20 s, 72°C for 20 s and fluorescence measurement. Following PCR, a thermal melting profile was used for amplicon identification.

For both quantification methods, standard curves were generated from four-fold dilutions of the corresponding genomic DNA. Serial dilutions of plasmids containing the loci of interest were used as standards for cDNA quantification. Each reaction was performed at least in duplicate. The second-derivative maximum method was used to quantify sequences, as described in the Light Cycler Software. For BrdU-labeled NS quantification, we used primer pairs binding to mitochondrial DNA to normalize enrichment in each S-phase fraction (Mitochondrial DNA), an early-replicated region (Early timing control) for the control of cell sorting, a genomic position located 5 kb downstream from insertion site 2 (Both), amplifying both alleles, for the control of accurate qPCR measurements and the studied regions in the transgenes (Figures 4, 5, S1 to S3 and Table S3). Specific primer pairs amplifying in the transgene (With) or the site of insertion in the wt allele (Without) were used (table S3). In the case of clones with one or two autonomous replicons associated with the GFP reporter construct inserted on the other chromosome, we used a primer pair amplifying the GFP reporter construct to quantify the wt chromosome (Without on GFP reporter at site 2, Table S3 and Figure S2C and E) and a primer pair amplifying insertion site 2 to quantify the modified shifting chromosome (With at insertion site 2, Table S3 and Figure S2C and E).

Based on both the one-hour BrdU pulse labeling used and the mean rate of fork progression, estimated at 1.5 kb/min in DT40 cells (Maya-Mendoza et al., 2007), the resolution of our RT experiments cannot exceed a 90 kb window. Thus, for the analysis of RT within our 30 kb genomic region targeted with one, two or three constructs, we systematically used the With and Without primer pairs in central position 2. For the quantitative analyses, we analyzed independent clones. In figure 2, *N*=10 for (i) from (Hassan-Zadeh et al., 2012), Figure 8 and (Valton et al., 2014), Figure S10, *N*=8 for (ii) from (Hassan-Zadeh et al., 2012), Figure 6B and (Valton et al., 2014), Figure S9, *N*=6 for (iii) from (Hassan-Zadeh et al., 2012), Figure 4A and this article Figure S1A and *N*=8 for (iv) from (Hassan-Zadeh et al., 2012), Figure 6A and this article, Figure S1B and S2B and C. In figure 5C, *N*=8 for (i) the same clones as (i) in Figure 2, *N*=5 for (ii) from Figure S3A-B, *N*=3 for (iii) from Figure S2 D-E.

For ChIp DNA and MNAse-digested DNA quantifications, we used primer pairs amplifying the wt allele at insertion sites 1, 2 and 3 (Wt allele insertion site 1, 2 or 3), the condensed region of the endogenous *β-globin* locus, as a control for the heterochromatin state (cond 1 and 2), the *BU1A* promoter (*BU1A* promoter) or the *MED14* promoter (*MED14* promoter), as a control for the active chromatin state, at a genomic position located 5 kb downstream from insertion site 2 (insertion site +5 kb), amplifying both alleles and the inserted transgenes, at the positions studied (Figure 3, 7 and Table S3).

For the quantification of MNAse-digested DNA, data are presented as total chromatin input relative to condensed genomic regions (cond1 and cond2). For each cell line, the mean values obtained for the two condensed genomic regions for each digestion were arbitrarily set to 1 and used to normalize independent clones for each transgenic line separately and between all transgenic lines.

For ChIp DNA quantifications, data are presented as Ip/Input signal relative to the signal for the *BU1A* promoter used for normalization between transgenic lines.

For cDNA quantification, we used primer pairs amplifying the second exon of the *BU1A* gene as a control for transcribed genes, the junction of exons 3-4 of the *MED14* gene and the inserted transgenes, at the studied positions (Figure 3 and Table S3).

