## Supplementary Materials for "Strong replicators associated with open chromatin are sufficient to establish an early replicating domain"

**Supplemental Information**

**Figure S1: Additional replication timing assays for transgenes shifting the timing of replication. Related to figure 2**

(A and B) RT profiles of each chromosomal allele determined after targeted transgene integration. Three different PCR primer sets were used to investigate the timing of replication at the integration site, for the modified allele (With), the endogenous wt allele (Without), or both alleles (Both). The endogenous *β-globin* locus was analyzed as an early-replicated control. Differences in -ΔL+ΔE and Δslope values calculated at the target site following transgene integration are indicated. Error bars correspond to the standard deviation for qPCR duplicates. (A) Analysis of five *BsR* clones described in Figure 2. (B) Analysis of two 2xFIV+*BsR* clones described in Figure 1. The black vertical bars represent the precise insertion position.

**Figure S2: One advanced replicon inserted at site 1 or at site 3 directs a shift to earlier replication independently of the presence of a *GFP* reporter construct inserted at site 2. Related to figures 2 and 5**

(A-E) RT profiles of each chromosomal allele were determined following targeted transgene integration. Three different PCR primer sets were used to investigate the timing of replication at the integration site, for the modified allele (With), the endogenous wt allele (Without), or both alleles (Both). The endogenous *β-globin* locus was analyzed as an early-replicated control. Differences in -ΔL+ΔE and Δslope values calculated at the target site following transgene integration are indicated. Error bars correspond to the standard deviation for qPCR duplicates. (A) Analysis of one clone containing one *GFP* reporter construct composed of the *GFP* reporter gene under the control of the $\beta^A$-*globin* promoter ($\beta^A$ pro) and linked to a 1.6 kb fragment of human chromosome 7 (h.K7) inserted at site 2. (B-C) Analysis of clones containing one autonomous replicon described in Figure 2 (2xFIV+*BsR*) inserted at site 1 and one *GFP* reporter construct inserted at site 2 on the same chromosome (B) or on the other chromosome (C). (D-E) Analysis of clones containing one autonomous replicon (2xFIV+*PuroR*) inserted at site 3 and one *GFP* reporter construct inserted at site 2 on the same chromosome (D) or on the other chromosome (E).

**Figure S3: Two advanced replicons inserted at sites 1 and 3 form an early-replicated domain. Related to figure 5**

(A and B) RT profiles of each chromosomal allele were determined following targeted transgene integration. Three different PCR primer sets were used to investigate the timing of replication at the integration site for the modified allele (With), the endogenous wt allele (Without), or both alleles (Both). The endogenous *β-globin* locus was analyzed as an early-replicated control. Differences in -ΔL+ΔE and Δslope values calculated at the target site following transgene integration are indicated.

Error bars correspond to the standard deviation for qPCR duplicates. Analysis of two 1+2+3 (A) and three 1+3 (B) clones described in Figure 5B.

**Figure S4: Validation of MNase digestion patterns before and after size selection. Related to Figures 3 and 7.**

Chromatin was extracted from two clones for each construct or combination (2xFIV-*BsR*, *BsR*, 2xFIV+*BsR*, 1+2+3, 1+3, described in Figure 2 and 5B) and partially digested with exponentially increasing concentrations of micrococcal nuclease (MNAse; 2.5, 10, 40 and 160 U/mL). After purification, DNA molecules were subjected to a size selection process that removed most DNA molecules over 1000 bp. The four digested DNA samples obtained for each clone were subjected to electrophoresis in a 1% w/v agarose gel before and after size selection and stained with SYBR safe. The DNA size marker was a commercial 1 kb plus ladder.

**Figure S5: PCR validation of clones selected for homologous recombination.**

(A-H) Schematic diagrams showing genomic region containing a site-specific integrated construct. The 5′ and 3′ arms of the targeted vector are shown as black boxes. Arrows #1 and #2 represent primer sets used for the analysis of correct integration of the constructs by homologous recombination. Other arrows represent primer sets used for the analysis of the correct excision of the *BsR* gene at site 2 (#3 and #4) or at site 3 (#5 and #6). PCR products were subjected to electrophoresis in a 1-1.5% w/v agarose gel and stained with SYBR safe. The DNA size marker was a commercial 1 kb plus DNA ladder (M). Lanes marked with a red star correspond to clones selected for further analysis. Insertion site 1, containing the *BsR* gene cassette (A), loxP_RE element (B) or the 2xFIV+*BsR* construct (C), is shown. (D) Insertion site 2 containing the *GFP* reporter construct is shown. (E) Insertion site 3, containing the 2xFIV+*PuroR* construct (E) is shown. The two genomic regions containing the 2xFIV+*BsR* construct inserted at the late1 (G) or late2 (H) site are shown. The sizes of the different PCR products obtained after amplification with primers #1 and #2, to check for correct integration, are 2.4 kb (A), 2.6 kb (B), 2.3 kb (C), 2.6 kb (D left gel), 2.3 kb (E), 2.6 kb (F) and 2.2 kb (G). The sizes of the different PCR products for the screening of clones correctly recombined for the *BsR* gene cassette after amplification with primers #3 and #4 are 2.1 kb (D, insertion site 2, right gel). The clone containing the *GFP* reporter construct without the *BsR* cassette at site 2 (D, 1-*BsR*) was used for further insertions at site 1 (B, 1 and C, 3-6) or at site 3 (E, 1-4). The clones containing the *GFP* reporter construct without the *BsR* cassette at site 2 and one autonomous replicon at site 1 (C, 3-4-5) were used for further insertions at site 3 (E 5-9). The clones containing the *GFP* reporter construct without the *BsR* cassette at site 2 and one loxP_RE at site 1 (B, 2) were used for further insertions of an autonomous replicon at site 3 (E, 10,11,12).

**Figure S6: Validation of transgenes integration into the same chromosome.**

(A) The insertion of transgenes into the same or a different chromosome was analyzed by long-range PCR, with a primer set composed of an upstream primer binding within one construct and a downstream primer binding to the other construct. The amplicons generated after PCR amplification to test for insertion into the same chromosome are represented by red arrows (LR1+2, LR2+3, LR1+3 and LR 1(loxP_RE)+3). The absence of an amplicon indicates that the two constructs were inserted into distinct chromosomes. The quality of the DNA was checked with primer sets amplifying, on the two chromosomes, a genomic region located between the two insertion sites (black arrows, LR 1+2 ctrl, LR 2+3 ctrl). (B) PCR products were subjected to electrophoresis in a 0.8% w/v agarose gel and stained with SYBR safe. The DNA size marker used was a commercial 20 kb plus ladder. 1+2 clonal lines #3 and #4 were used to generate 1+3 clonal lines #1 and #2 with the *GFP* reporter construct inserted at site 2 on the same chromosome and 1+2 clonal line #2 was used to generate 1+3 clonal lines #3-5 with the *GFP* reporter construct inserted at site 2 on the other chromosome. Clone 1 with the 2xFIV construct inserted at site 1, as previously obtained (Hassan-Zadeh 2012, Figure 6B), was used to generate 1+3 clonal line # 6-10. Clone #1 with the *GFP* reporter construct inserted at site 2 (Figure S5D) was used to generate one clone with the loxP_RE element inserted at site 1, which itself generated 1(loxP_RE)+3 clonal lines #1, 2 and 3 with the *GFP* reporter construct inserted at site 2 on the other chromosome.

### Tables

**Table S1: Cell lines summary table**

**Table S2: Transgene copy number determination in cell lines**

The table shows the qPCR results obtained with genomic DNA extracted from the clones selected for the experiments. For each clone, 2 ng of genomic DNA was amplified with a primer set amplifying a sequence within the construct (With) and another primer set amplifying a sequence 5 kb downstream from the insertion site for both alleles (Both). The ratio of the amounts of DNA obtained with the With and Both primer sets was used to determine transgene copy number in all cell lines.

**Table S3: Primer sets used for quantitative PCR**

Figure S1

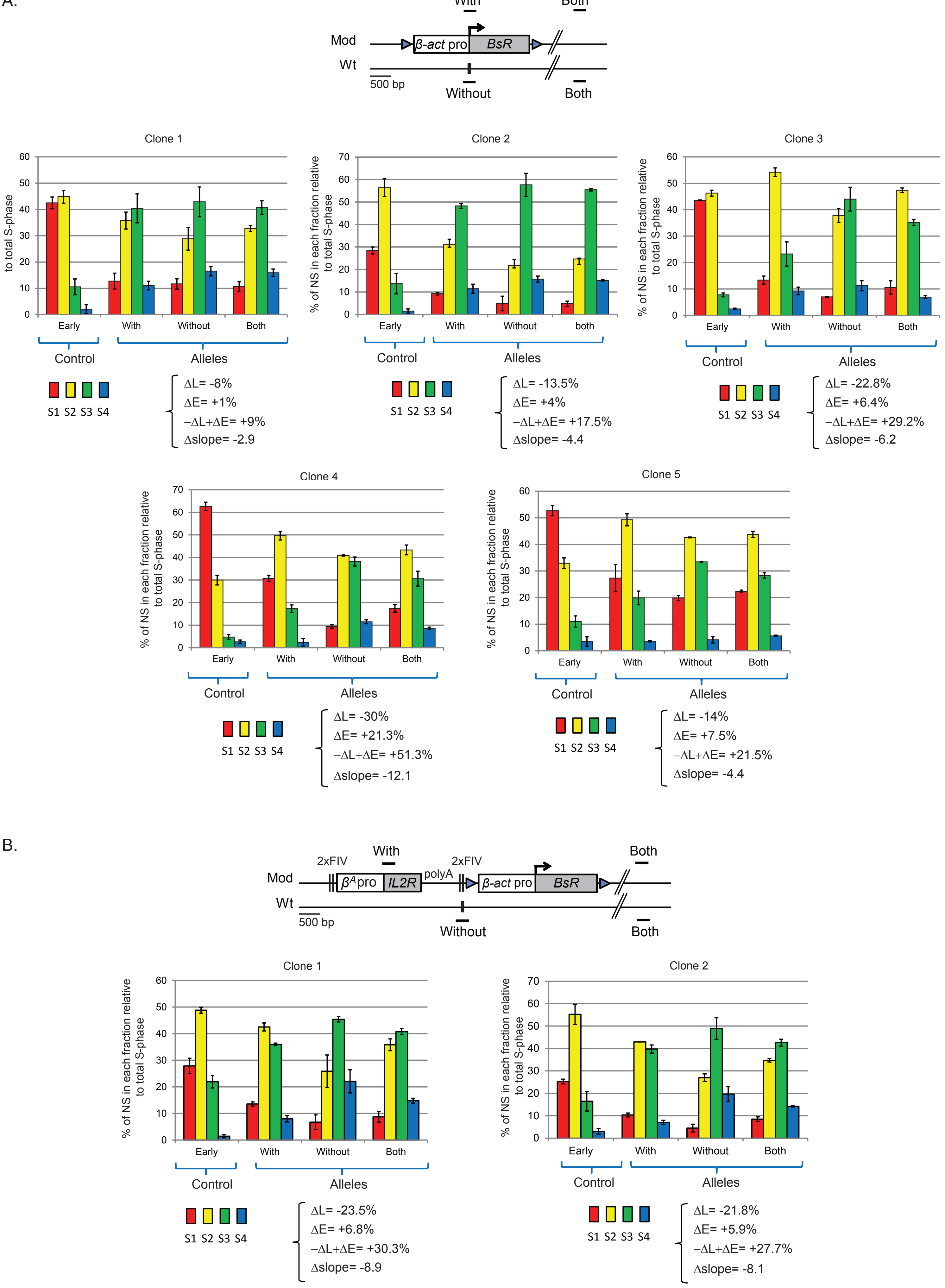
A.
With
Both
Mod
β-act pro
BsR
Wt
500 bp
Without
Both
Clone 1
Clone 2
Clone 3
% of NS in each fraction relative to total S-phase
Early
With
Without
Both
both
Control
Alleles
S1 S2 S3 S4
ΔL= -8%
ΔE= +1%
−ΔL+ΔE= +9%
Δslope= -2.9
ΔL= -13.5%
ΔE= +4%
−ΔL+ΔE= +17.5%
Δslope= -4.4
ΔL= -22.8%
ΔE= +6.4%
−ΔL+ΔE= +29.2%
Δslope= -6.2
Clone 4
Clone 5
ΔL= -30%
ΔE= +21.3%
−ΔL+ΔE= +51.3%
Δslope= -12.1
ΔL= -14%
ΔE= +7.5%
−ΔL+ΔE= +21.5%
Δslope= -4.4
B.
2xFIV
With
2xFIV
Both
polyA
βApro
IL2R
β-act pro
BsR
Mod
Wt
500 bp
Without
Both
Clone 1
Clone 2
ΔL= -23.5%
ΔE= +6.8%
−ΔL+ΔE= +30.3%
Δslope= -8.9
ΔL= -21.8%
ΔE= +5.9%
−ΔL+ΔE= +27.7%
Δslope= -8.1

Figure S2

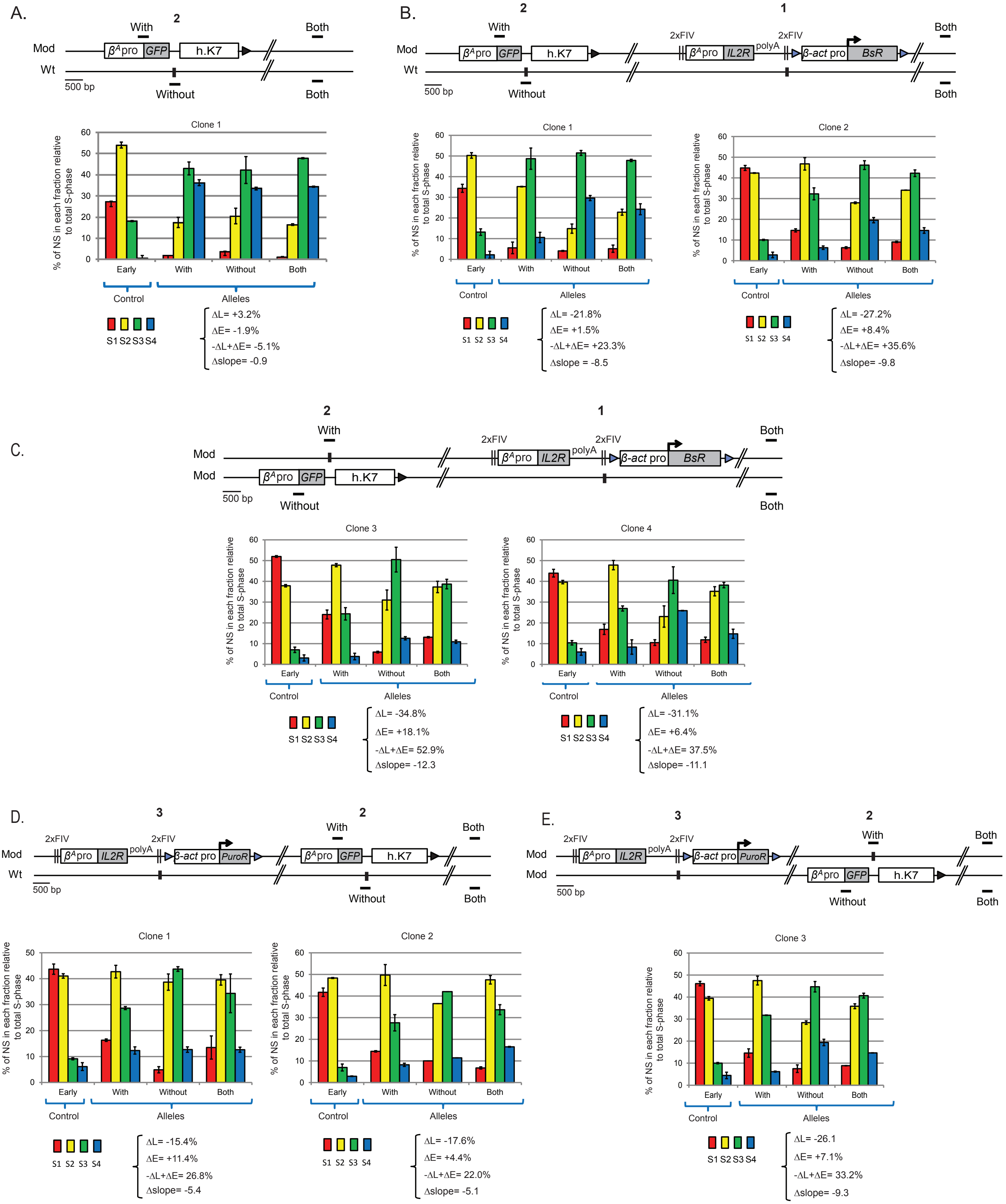
A.
Mod
Wt
With
Without
Both
2
βApro GFP
h.K7
500 bp
Clone 1
% of NS in each fraction relative to total S-phase
Early
With
Without
Both
Control
Alleles
S1 S2 S3 S4
ΔL= +3.2%
ΔE= -1.9%
-ΔL+ΔE= -5.1%
Δslope= -0.9
B.
2
1
2xFIV
polyA
βApro IL2R
β-act pro BsR
Clone 1
ΔL= -21.8%
ΔE= +1.5%
-ΔL+ΔE= +23.3%
Δslope = -8.5
Clone 2
ΔL= -27.2%
ΔE= +8.4%
-ΔL+ΔE= +35.6%
Δslope= -9.8
C.
Clone 3
ΔL= -34.8%
ΔE= +18.1%
-ΔL+ΔE= 52.9%
Δslope= -12.3
Clone 4
ΔL= -31.1%
ΔE= +6.4%
-ΔL+ΔE= 37.5%
Δslope= -11.1
D.
3
β-act pro PuroR
Clone 1
ΔL= -15.4%
ΔE= +11.4%
-ΔL+ΔE= 26.8%
Δslope= -5.4
Clone 2
ΔL= -17.6%
ΔE= +4.4%
-ΔL+ΔE= 22.0%
Δslope= -5.1
E.
Clone 3
ΔL= -26.1
ΔE= +7.1%
-ΔL+ΔE= 33.2%
Δslope= -9.3

Figure S3

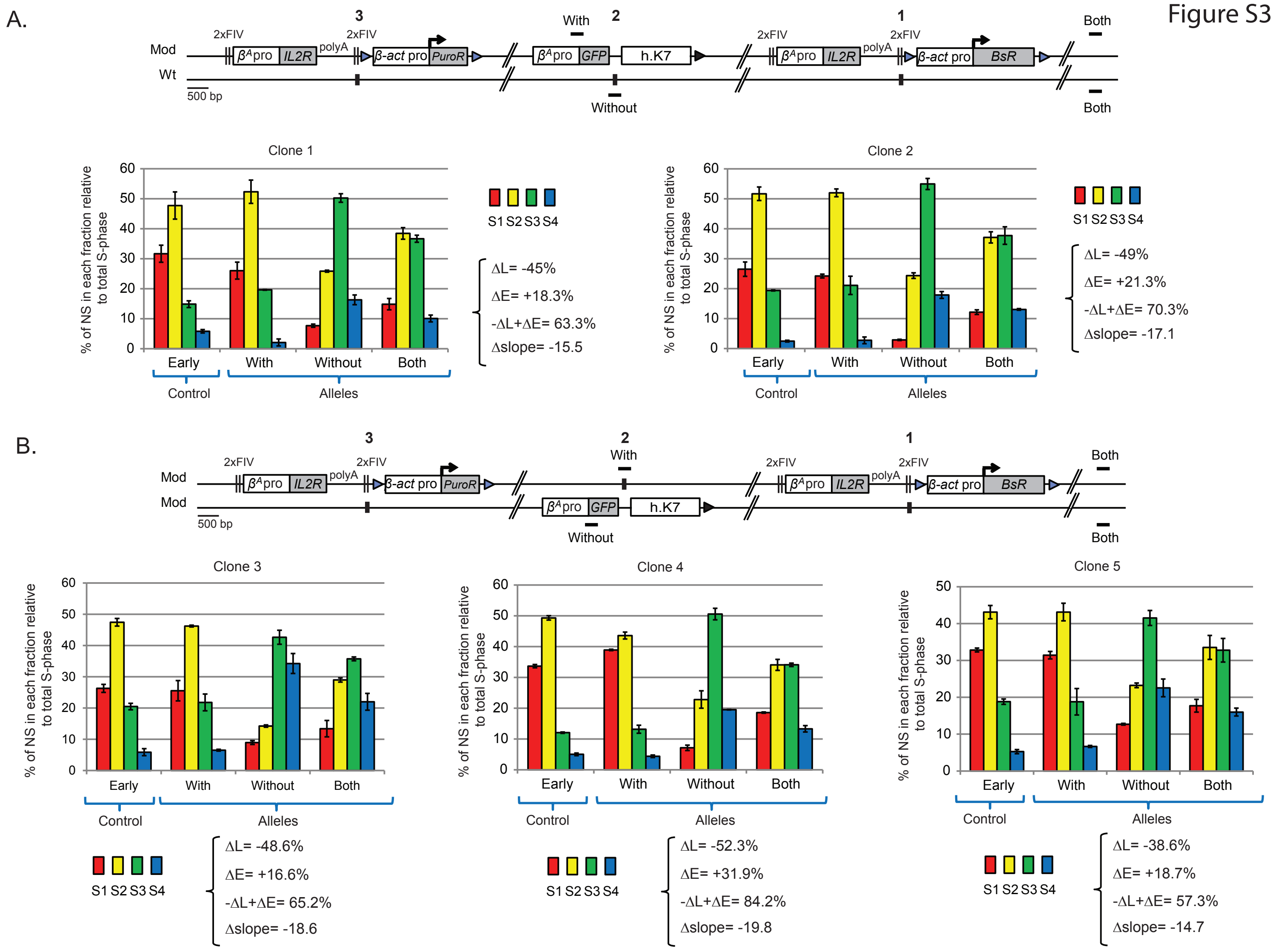
A.
Mod
Wt
3
2
1
With
Without
Both
2xFIV
polyA
βApro
IL2R
β-act pro
PuroR
GFP
h.K7
BsR
500 bp
Clone 1
Clone 2
% of NS in each fraction relative to total S-phase
Early
With
Without
Both
Control
Alleles
S1 S2 S3 S4
ΔL= -45%
ΔE= +18.3%
-ΔL+ΔE= 63.3%
Δslope= -15.5
ΔL= -49%
ΔE= +21.3%
-ΔL+ΔE= 70.3%
Δslope= -17.1
B.
Mod
Mod
Clone 3
Clone 4
Clone 5
ΔL= -48.6%
ΔE= +16.6%
-ΔL+ΔE= 65.2%
Δslope= -18.6
ΔL= -52.3%
ΔE= +31.9%
-ΔL+ΔE= 84.2%
Δslope= -19.8
ΔL= -38.6%
ΔE= +18.7%
-ΔL+ΔE= 57.3%
Δslope= -14.7

Figure S4

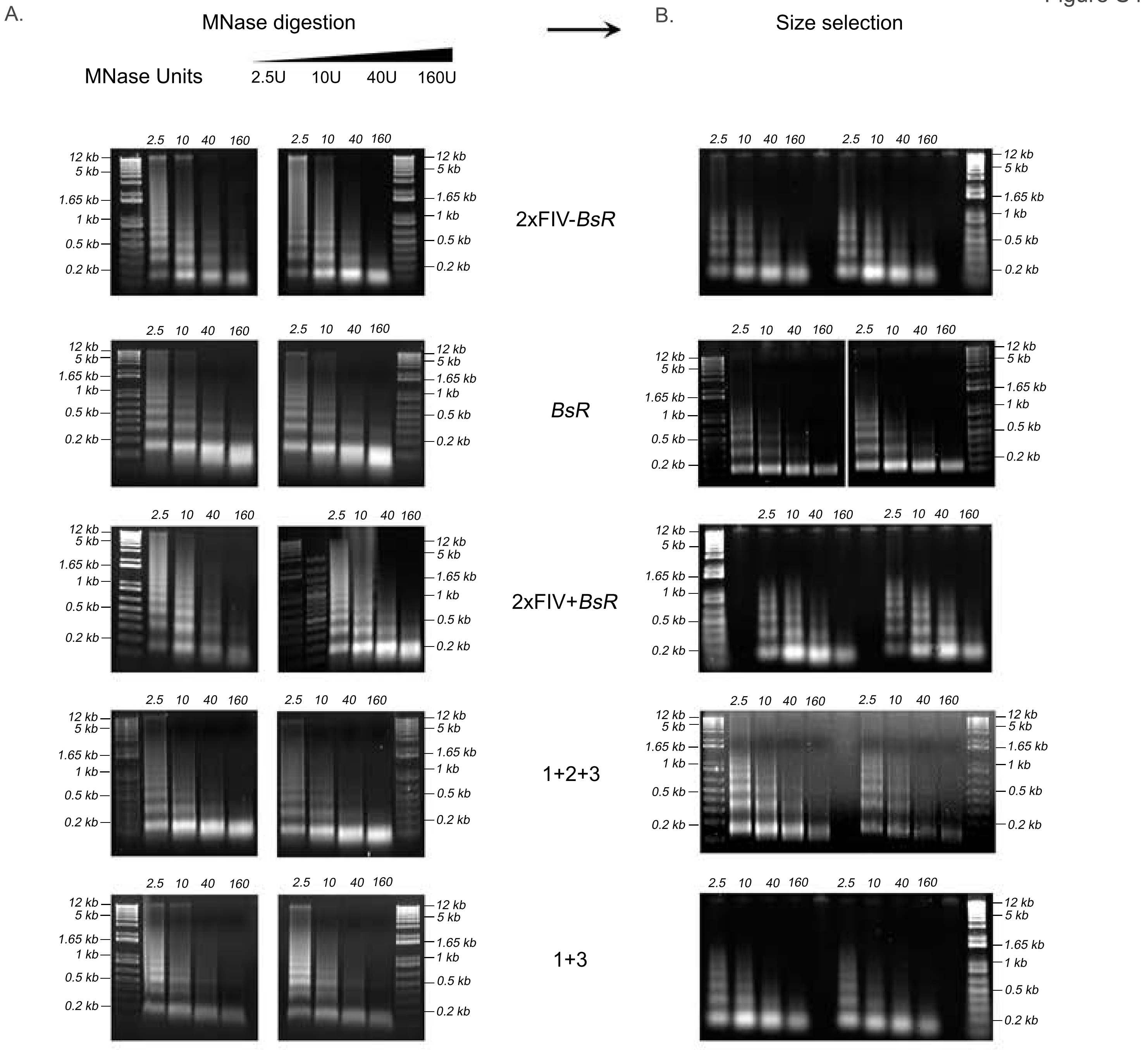

A.
MNase digestion
MNase Units
2.5U
10U
40U
160U
B.
Size selection
2.5
10
40
160
12 kb
5 kb
1.65 kb
1 kb
0.5 kb
0.2 kb
2xFIV-*BsR*
*BsR*
2xFIV+*BsR*
1+2+3
1+3

Figure S5

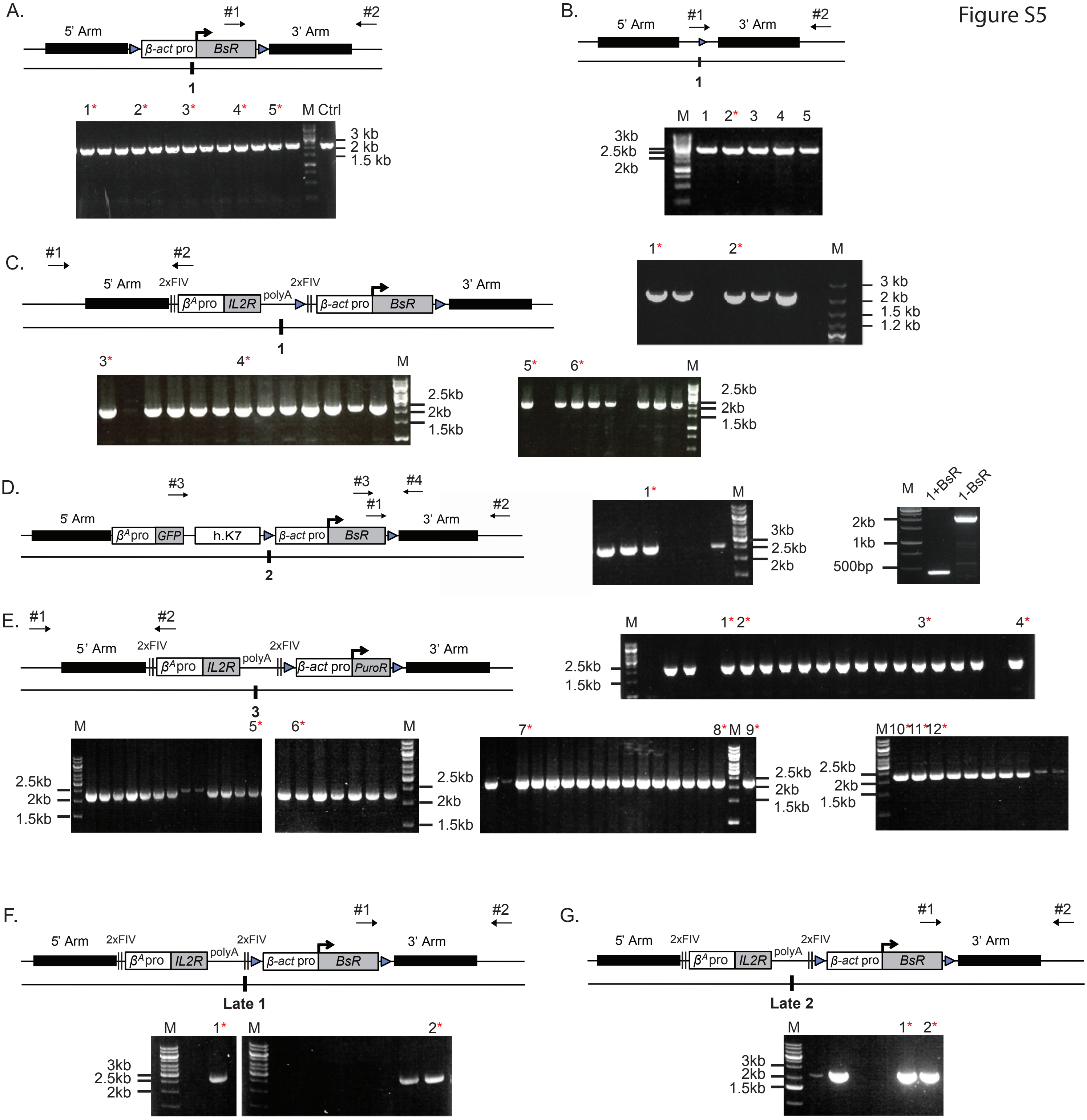
A.
5' Arm
β-act pro
BsR
3' Arm
#1
#2
1
1* 2* 3* 4* 5* M Ctrl
3 kb
2 kb
1.5 kb
B.
5' Arm
3' Arm
#1
#2
1
M 1 2* 3 4 5
3kb
2.5kb
2kb
1* 2* M
3 kb
2 kb
1.5 kb
1.2 kb
C.
#1
#2
5' Arm
2xFIV
βApro
IL2R
polyA
2xFIV
β-act pro
BsR
3' Arm
1
3* 4* M
2.5kb
2kb
1.5kb
5* 6* M
2.5kb
2kb
1.5kb
D.
#3
#3
#4
#1
#2
5' Arm
βApro
GFP
h.K7
β-act pro
BsR
3' Arm
2
1* M
3kb
2.5kb
2kb
M 1+BsR 1-BsR
2kb
1kb
500bp
E.
#1
#2
5' Arm
2xFIV
βApro
IL2R
polyA
2xFIV
β-act pro
PuroR
3' Arm
3
M 1* 2* 3* 4*
2.5kb
1.5kb
M 5*
6* M
2.5kb
2kb
1.5kb
7* 8* M 9*
M 10* 11* 12*
F.
#1
#2
5' Arm
2xFIV
βApro
IL2R
polyA
2xFIV
β-act pro
BsR
3' Arm
Late 1
M 1* M 2*
3kb
2.5kb
2kb
G.
#1
#2
5' Arm
2xFIV
βApro
IL2R
polyA
2xFIV
β-act pro
BsR
3' Arm
Late 2
M 1* 2*
3kb
2kb
1.5kb

Figure S6

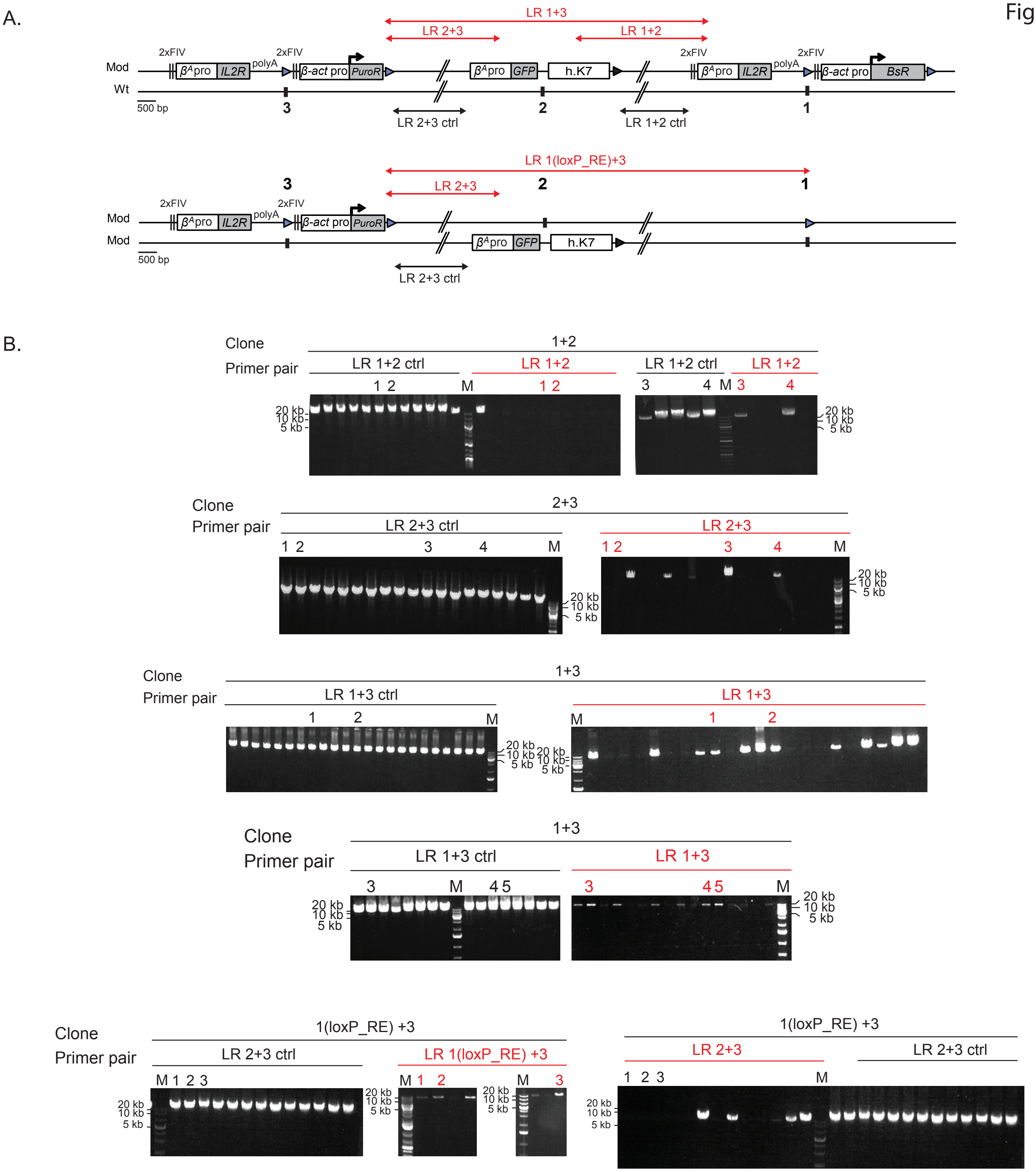
A.
LR 1+3
LR 2+3
LR 1+2
2xFIV
polyA
Mod
Wt
500 bp
β^A pro
IL2R
β-act pro
PuroR
GFP
h.K7
BsR
3
2
1
LR 2+3 ctrl
LR 1+2 ctrl
LR 1(loxP_RE)+3
B.
Clone
Primer pair
1+2
2+3
1+3
1(loxP_RE) +3
LR 1+2 ctrl
LR 1+2
LR 2+3 ctrl
LR 2+3
LR 1+3 ctrl
LR 1+3
LR 1(loxP_RE) +3
M
20 kb
10 kb
5 kb

<table>
<tr><th>Site of insertion</th><th>Number of clones<br>and source</th><th>RT shift<br>Median values</th></tr>
<tr><td>Mid-Late (1)</td><td>Total= 44</td><td></td></tr>
<tr><td>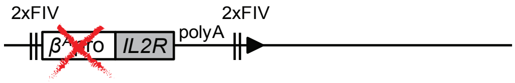</td><td>10 { 2 Hassan-Zadeh et al. 2012 (Figure 8)<br>8 Valton et al. 2014 (Figure 8C, S10)</td><td>-ΔL+ΔE= +6.5%<br>Δslope= -1.6</td></tr>
<tr><td>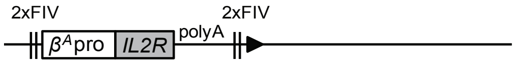</td><td>8 { 2 Hassan-Zadeh et al. 2012 (Figure 6B)<br>6 Valton et al. 2014 (Figure 8C, S9)</td><td>-ΔL+ΔE= +20.2%<br>Δslope= -5.6</td></tr>
<tr><td>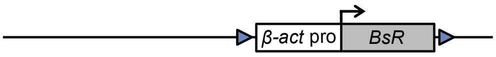</td><td>6 { 1 Hassan-Zadeh et al. 2012 (Figure 4A)<br>5 This study (Figure S1A)</td><td>-ΔL+ΔE= +19.6%<br>Δslope= -5.3</td></tr>
<tr><td>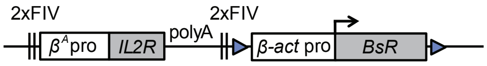</td><td>8 { 2 Hassan-Zadeh et al. 2012 (Figure 6A)<br>6 This study (Figure S1B, S2B, C)</td><td>-ΔL+ΔE= +36.6%<br>Δslope= -10.1</td></tr>
<tr><td>Mid-Late (3)</td><td></td><td></td></tr>
<tr><td>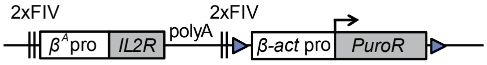</td><td>3 { This study (Figure S2D, E)</td><td>-ΔL+ΔE= +26.8%<br>Δslope= -5.4</td></tr>
<tr><td>Mid-Late (1+3 and 1+2+3)</td><td></td><td></td></tr>
<tr><td>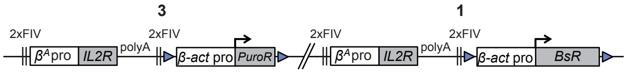</td><td>5 { This study (Figure S3)</td><td>-ΔL+ΔE= +65.2%<br>Δslope= -17.1</td></tr>
<tr><td>Late (1)</td><td></td><td></td></tr>
<tr><td>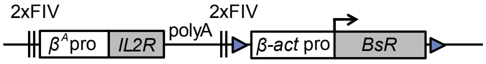</td><td>2 { This study (Figure 4B)</td><td>-ΔL+ΔE=<br>+29.3% / +51.7%<br>Δslope= -12.9/-16.7</td></tr>
<tr><td>Late (2)</td><td></td><td></td></tr>
<tr><td>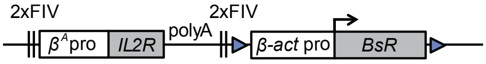</td><td>2 { This study (Figure 4C)</td><td>-ΔL+ΔE=<br>+16.1% /+19.1%<br>Δslope= -8.2/-10.6</td></tr>
</table>

Table S2

| | Both concentration | With concentration | Ratio With/Both |
|---|---|---|---|
| Insertions at site mid-1 | | | |
| *BsR* (1) | 0.97 | 0.45 | 0.47 |
| *BsR* (2) | 0.78 | 0.32 | 0.41 |
| *BsR* (3) | 0.63 | 0.37 | 0.59 |
| *BsR* (4) | 0.97 | 0.62 | 0.63 |
| *BsR* (5) | 0.93 | 0.41 | 0.44 |
| 2xFIV +*BsR* (1) | 1.34 | 0.71 | 0.53 |
| 2xFIV +*BsR* (2) | 1.29 | 0.65 | 0.5 |
| $\beta^A$*GFP* h.K7+2xFIV +*BsR* (1) | 0.81 | 0.45 | 0.55 |
| $\beta^A$*GFP* h.K7+2xFIV +*BsR* (2) | 0.84 | 0.43 | 0.51 |
| $\beta^A$*GFP* h.K7/2xFIV +*BsR* (3) | 0.74 | 0.4 | 0.54 |
| $\beta^A$*GFP* h.K7/2xFIV +*BsR* (4) | 0.91 | 0.45 | 0.49 |
| LoxP_RE (1) | 0.58 | 0.28 | 0.48 |
| Insertions at site mid-2 | | | |
| $\beta^A$*GFP* h.K7 (1) | 1.12 | 0.54 | 0.49 |
| Insertions at site mid-3 | | | |
| 2xFIV+ *PuroR* + $\beta^A$*GFP* h.K7 (1) | 1.04 | 0.57 | 0.55 |
| 2xFIV+ *PuroR* + $\beta^A$*GFP* h.K7 (2) | 1.59 | 0.59 | 0.37 |
| 2xFIV+ *PuroR* / $\beta^A$*GFP* h.K7 (3) | 1.08 | 0.43 | 0.4 |
| Insertions at site mid-1 and mid-3 | | | |
| 2xFIV+ *PuroR* + $\beta^A$*GFP* h.K7+2xFIV +*BsR* (1) | 0.56 | 0.73 | 1.29 |
| 2xFIV+ *PuroR* + $\beta^A$*GFP* h.K7+2xFIV +*BsR* (2) | 0.54 | 0.58 | 1.07 |
| 2xFIV+ *PuroR* / $\beta^A$*GFP* h.K7/2xFIV +*BsR* (3) | 1.15 | 1.33 | 1.16 |
| 2xFIV+ *PuroR* / $\beta^A$*GFP* h.K7/2xFIV +*BsR* (4) | 0.87 | 0.99 | 1.13 |
| 2xFIV+ *PuroR* / $\beta^A$*GFP* h.K7/2xFIV +*BsR* (5) | 0.83 | 0.94 | 1.13 |
| 2xFIV+ *PuroR* +LoxP_RE (1) | 3.34 | 2.21 | 0.66 |
| 2xFIV+ *PuroR* +LoxP_RE (2) | 2.19 | 1.39 | 0.63 |
| 2xFIV+ *PuroR* +LoxP_RE (3) | 2.12 | 1.13 | 0.53 |
| Insertions at site late-1 | | | |
| 2xFIV +*BsR* (1) | 2.15 | 0.96 | 0.45 |
| 2xFIV +*BsR* (2) | 1.71 | 0.75 | 0.44 |
| Insertions at site late-2 | | | |
| 2xFIV +*BsR* (1) | 1.6 | 0.59 | 0.37 |
| 2xFIV +*BsR* (2) | 1.37 | 0.5 | 0.36 |

Table S3

| | Figures | Forward primer sequence | Reverse primer sequence | Genomic position (Build Dec 2015) |
|---|---|---|---|---|
| Replication timing analysis | | | | |
| With on *GFP* reporter | 5,S2A,B,D, S3A | GGAATTCGATAGCTTGGCGGC | GCTGAACTTGTGGCCGTTTAC | |
| With on insertion site 2 | 5, S2C,E, SB | | | |
| With on 2xFIV+*BsR or* | 4, S1B, table1 | GGGGACTGCTCACGTTCATCA | AATGTGGCGTGTGGGATCTC | |
| With on *BsR* | S1A, table1 | TGCAGAAATCGGAGGAAGAAGA | GAATTGCCGCTCCCACATGA | |
| Without or Wt allele insertion site 1 | 3, 7, S1 | CAGGACAGCAGGTATTCACA | GGCCTGAACACTGTGTCAAT | chr1:72565497+72565651 |
| Without or Wt allele insertion site 2 | 5, 7, S2A,B,D, S3A | GTAATGAAATTCAGCAATGACAGGC | TCCTATCTGTTCAAATGTGCATCAG | chr1:72548543+72548676 |
| Without on GFP reporter at site 2 | 5, S2C,E, S3B | | | |
| Wt allele insertion site 3 | 7 | TGGTACAGGCTGAGGACACC | TGATGACTGCAGCTTCCTTCT | chr1:72535996+72536105 |
| Without or Wt allele insertion | 4 | CCCTTGAATCAGACCCTTGA | CCCTCCTTTCTCCATAAAAACA | chr1:70523547+70523674 |
| Without or Wt allele insertion | 4 | TTTACACTACTCCCACCCCTCG | TTGACCATATGCCACCAACACC | chr1:177936339+177936438 |
| 3' proximal early domain | | | | |
| +400kb | 5 | | | |
| +500kb | 5 | | | |
| Controls | | | | |
| Both or insertion site 1 + 5kb | 3, 5, 7, S1, S2, S3, table1 | TCCATACAGCCACAACAGCA | TGTGGAAGAGTTTCAGTCCAGG | chr1:72570952+72571067 |
| Both or insertion late 1 - 4.8kb | 4 | TGTACTTCTCTGTGGACATGCA | TGGCACAGAGGACAGGTAAGA | chr1:70518727+70518810 |
| Both or insertion late 2 - 3.6kb | 4 | CAAGGTTTCCACCCCTAAAGA | TGATGGATGTGGGAAGAGAAA | chr1:177932452+ 177932533 |
| Early timing control | 4,5, S1, S2, S3 | GACGGTCAGGTTTGCCAAAG | TCCTGAGGATACGTTTTTCAG | chr1:194563998+194564262 |
| Mitochondrial DNA | 4, 5, S1, S2, | CATCCCATGCATAACTCCTG | GTAGTCCAGGCTTCACTTGA | chrM:541+731 |
| ChIps, Chromatin accessibility and RNA quantification | | | | |
| 5' 2xFIV site 1 | 3, 7 | GGGCTATTCAGCTTGTCTAG | GCCACCTCAACTTTTGTATAC | |
| 5' 2xFIV site 3 | 7 | TTATGCTGGCAGGACTGAGA | GTGGGCAGAGGAAAGCGAT | |
| $\beta^A$ promoter 1 | 3 | TCCGGAGATGCAGCCAATTG | CCGCCAAGCTATCGAATTCCTG | |
| $\beta^A$ promoter 2 | 3, 7 | GGGAGCAAGAGCCCAGAC | GTGAGCAGTCCCCACATCAG | |
| $\beta^A$ promoter 3 | 3, 7 | GGGACTGCTCACGTTCATCA | AATGTGGCGTGTGGGATCTC | |
| *IL2R* gene | 3, 7 | CTACACAGAGGTCCTGCTG | GTGAAGAGAAGCCTCAGGCA | |
| 3' 2xFIV site 1 | 3 | TGCATTCTAGTTGTGGTTTGTCC | ACCGTCGACCAACTTTGTATAGA | |
| 3' 2xFIV site 1 and 3 | 7 | AAGCTTGGATCCCCTACCGT | GAGAGTGAAGCAGAACGTGGG | |
| 5' β-actin promoter site 1 | 3 | | | |
| *β-actin* promoter site 1 | 3, 7 | TGCAGAAATCGGAGGAAGAAGA | GAATTGCCGCTCCCACATGA | |
| *BsR* gene | 3, 7 | CGGCAGTACATATTGAAGCGT | CCCTACACATACCACAAGGA | |
| *PuroR* gene | 7 | ACGACCTTCCATGACCGAGT | AGTTCTTGCAGCTCGGTGAC | |
| $\beta^A$ *-GFP* | 7, table1 | GGAATTCGATAGCTTGGCGGC | GCTGAACTTGTGGCCGTTTAC | |
| *GFP* gene | 7 | GCCCGACAACCACTACCTGAG | GCTTTACTTGTACAGCTCGTCCA | |
| h.K7 | 7 | AAGTTTATCATTGTGTGGCAGTCA | GTTCTGGTTGGGTTGTATGTCAC | |
| cond 1 | 3, 7 | CATCTGTGCTCTGGGTCCA | AAGGAGTGGAAGGAACGCATC | chr1:194546368+194546497 |
| cond 2 | 3, 7 | TTGGTGCAGTGCCTCAGATAG | ATGTCGCTTGTCACGATGGAT | chr1:194546457+194546563 |
| *MED14* promoter | 3, 7 | GGATTCACACTGTTCCCCTCC | TGCATGTTTCTCTCATCCGAAGT | chr1:112227330+112227461 |
| *BU1A* promoter | 3 | CTCTGTAGCCAGATCGTCTTCTC | GTGTCAGCTCATCTAGGCAAATC | chr1:91922377+91922546 |
| *BU1A* gene | 3 | AATGTCCCCAAAATGAGCTG | CCTCTTTTTCCACCCTCCTC | chr1:91923373+91923517 |
| *Med14* gene | 3 | TGGGCTAATAATGCTGGAAAGGT | TAGAGAAGCCAGACGATCAGCA | |
| Long range amplifications | | | | |
| LR 1+2 | S6 | AAGGGTCAGCTTTCGTGATAATCTGG | ACCTCTCTTGCATTCACAGTTCAACA | |
| LR 1+2 ctrl | S6 | GCAAGATGGGCAGAGCTGAGTTAACAAT | TGTCCTGTAAGTCCTGGCAAAACAAAG A | chr1:72548824+72565503 |
| LR 2+3 | S6 | GAGCGTATTACAATTCACTGGCCGTC | CTTGCTCACCATTTCCTGACCCTTG | |
| LR 2+3 ctrl | S6 | GCAGTATAACAAGCACGCCTGAAGTAAA | CAGTCTTATCCCACCCCTTCCTGATAG | chr1:72536307+72548324 |
| LR 1+3 | S6 | GAGCGTATTACAATTCACTGGCCGTC | GATCCCGTGCCACCTCAACTTTTGTAT | |
| LR 1(loxP_RE)+3 | S6 | GAGCGTATTACAATTCACTGGCCGTC | GACGTTGTGGCTGTTGTAGTTGTACTC | |
| Screening of targetted integration | | | | |
| 3'-screening-site 2 for *GFP* reporter | S5D | GCTCCAATTCGCCCTATAGTGA | GGCACTCCATTTCCATCTCCT | |
| 5'-screening-site 3 for 2xFIV+*PuroR* | S5E | GGACTGGCTAGGGAACAAGAG | TCTGCCTTCTCCCTGATAACG | |
| 5'-screening-site 1 for 2xFIV+*BsR* | S5C | GTGCAGCATCAGTGGATAAAGT | TCTGCCTTCTCCCTGATAACG | |
| 5'-screening-site1 for *BsR* | S5A | CCCCCTGAACCTGAAACATAA | CCACATGTTTATTGCATACGGC | |
| 3'-screening-site1 for loxP_RE | S5B | CCAATTCGCCCTATAGTGAGTCG | ACGTAACAAATCTACAGGTCTTCG | |
| 3'-screening-site late-1 for | S5F | GAGCTCCAATTCGCCCTAT | ACTATTGTACCCCTCCCTGTTG | |
| 3'-screening-site late-2 for | S5G | GAGCTCCAATTCGCCCTAT | GGTCTGATCCCTATCTCACTGG | |
| Screening of site specific excision | | | | |
| Large 1+3 excision | 6 | GGTTCTGGTGCCCTCATTGAT | TGCTCTGCTAGAATTGCCTGT | chr1:72535815+72565623 |
| Ctrl | 6 | ACCCAAGGCAGGCTACAAAC | TGAGTTACTTTGGCATTACTTTTCATC | chr1:72565577+72567483 |
| Copy number quantification | | | | |
| LoxP_RE | table1 | GGAGGTCTTTGTTTTGCCAGGA | ACTAGTGGATCCCCTACCGTTC | |
